# A pan-plant protein complex map reveals deep conservation and novel assemblies

**DOI:** 10.1101/815837

**Authors:** Claire D. McWhite, Ophelia Papoulas, Kevin Drew, Rachael M. Cox, Viviana June, Oliver Xiaoou Dong, Taejoon Kwon, Cuihong Wan, Mari L. Salmi, Stanley J. Roux, Karen S. Browning, Z. Jeffrey Chen, Pamela C. Ronald, Edward M. Marcotte

**Author notes:** These authors contributed equally to this work (Claire D. McWhite, Ophelia Papoulas).

## Abstract

Plants are foundational to global ecological and economic systems, yet most plant proteins remain uncharacterized. Protein interaction networks often suggest protein functions and open new avenues to characterize genes and proteins. We therefore systematically determined protein complexes from 13 plant species of scientific and agricultural importance, greatly expanding the known repertoire of stable protein complexes in plants. Using co-fractionation mass spectrometry, we recovered known complexes, confirmed complexes predicted to occur in plants, and identified novel interactions conserved over 1.1 billion years of green plant evolution. Several novel complexes are involved in vernalization and pathogen defense, traits critical to agriculture. We also uncovered plant analogs of animal complexes with distinct molecular assemblies, including a megadalton-scale tRNA multi-synthetase complex. The resulting map offers the first cross-species view of conserved, stable protein assemblies shared across plant cells and provides a mechanistic, biochemical framework for interpreting plant genetics and mutant phenotypes.

## INTRODUCTION

Plants make up most of the planet’s biomass and sustain global economic and environmental systems. Despite increasing numbers of sequenced plant genomes, biochemical characterization of encoded plant proteins captures a relatively small portion of the expected breadth of biological functions. The best characterized dicotyledonous plant, *Arabidopsis thaliana*, encodes approximately 35,000 protein-coding genes, but the functions of the majority of these proteins remain uncharacterized, even by homology (Rhee and Mutwil, 2014). This trend is similar for *Oryza sativa* (rice), a critical food crop, and the best characterized monocotyledonous plant (Kawahara et al., 2013). In marked contrast to other model organisms, it is estimated that only 5% of *Arabidopsis* proteins and considerably fewer proteins in other plants have had biochemical activity, localization, and biological roles determined by direct experimentation (Niehaus et al., 2015; Rhee and Mutwil, 2014; Swarbreck et al., 2008). Extraordinary experimental efforts are needed to define the core expressed proteome and molecular machinery of plants.

The active multiprotein assemblies in plant cells have not yet been systematically defined, in stark contrast to recent progress in humans (Havugimana et al., 2012; Hein et al., 2015; Huttlin et al., 2015, 2017; Kirkwood et al., 2013; Kristensen et al., 2012; Malovannaya et al., 2011; Wan et al., 2015). Determining protein-protein interactions is a key step for discovering gene and protein function and frequently opens new avenues to study and manipulate critical cellular processes (Eisenberg et al., 2000; Hartwell et al., 1999; Schwikowski et al., 2000; Walhout and Vidal, 2001). In model organisms, such as yeast or *Drosophila*, systematic mapping of protein complexes led to critical functional insights, facilitated understanding of conserved and disease-related pathways (Vidal et al., 2011), and helped characterize uncharacterized proteins *via* guilt by association (Hu et al., 2009; Marcotte et al., 1999; Oliver, 2000). Revealing this class of biochemical information for plants will dramatically advance efforts in fundamental plant research, as well as guide practical applications such as improvements in crop yields, disease/stress resistance, and biofuel production.

Unfortunately, many of the techniques that have been used to discover protein complexes at scale in animals and yeast (e.g., high-throughput affinity purification (AP-MS)) are prohibitively expensive or even intractable at large-scale in plants due to issues of complex genomes, polyploidy, and transformation efficiency. Consequently, AP-MS experiments in plants have been limited to targeted protein families, primarily in *Arabidopsis* and rice (Bassel et al., 2012; Rohila et al., 2009; Van Leene et al., 2008, 2010, 2011; Zhang et al., 2010). Such approaches for studying protein-protein interactions are disproportionately difficult in less well-characterized plant species necessitating new strategies for comparative studies.

Co-fractionation mass spectrometry (CF-MS) is a high-throughput method to detect interacting proteins, applicable to any organism, without the need for antibodies or transgenic epitope tagging of individual proteins. CF-MS involves chromatographically separating a native protein extract, then identifying proteins in each biochemical fraction using mass spectrometry. Co-elution (co-fractionation) of proteins in a separation serves as evidence of physical association, which, when measured over multiple distinct separations is a rigorous signal of protein interaction (Wan et al., 2015). Observation of repeated co-elution in multiple different species reduces species-specific artifacts and adds power to discover conserved—and thus more likely functional—interactions. Furthermore, the use of machine learning methods allows for strong control over false discovery rates of protein interactions.

Several papers have classified candidate *Arabidopsis* protein complexes using native (non-denaturing) separations such as gels (Takabayashi et al., 2017) or co-fractionation (Aryal et al., 2017; Gilbert and Schulze, 2019; McBride et al., 2019). However, in a single or small set of experiments, two proteins can coincidentally co-elute without a physical association. By comparison, repeated co-elution using multiple distinct species, tissues, and biochemical separations provides increasing confidence in a protein interaction (Wan et al., 2015). As with genome-wide association studies (GWAS) or recombination mapping of a single mutation by phenotype, large amounts of data are required to build a statistically strong observation.

In this largest survey to date of expressed plant proteins as well as their physical assemblies, we collected a massive and diverse plant co-elution dataset to define high confidence protein-protein interactions in a statistical computational framework, recovering known complexes and identifying novel complexes conserved across plants for over a billion years. Multiple newly discovered complexes have direct relevance to agronomically important traits. Collectively, the resulting set of protein abundances and map of stable protein interactions will help interpret plant gene functions in a mechanistic, evolutionary, and biochemical framework.

## RESULTS AND DISCUSSION

### A massive data set of protein abundances and co-purification from 13 plant species

We generated a large, diverse, and representative proteomics dataset from 13 species spanning 1.1 billion years of green plant evolution (**Figure 1A**). Our compendium incorporates proteomic data from *Arabidopsis thaliana, Brassica oleracea* (broccoli), *Glycine max* (soy), *Cannabis sativa* (hemp), *Solanum lycopersicum* (tomato), *Chenopodium quinoa* (quinoa), *Zea mays* (maize), *Oryza sativa ssp. japonica* (rice), *Triticum aestivum* (wheat), *Cocos nucifera* (coconut), *Ceratopteris richardii* (fern), *Selaginella moellendorffii* (spikemoss), and the green algae *Chlamydomonas reinhardtii*. We chose crop and model plants important to the research community, including species that overcome specific technical challenges (e.g. high yields of nuclei from broccoli and coconut; embryonic tissue from wheat and hemp). We included primitive vascular plants (fern, spikemoss) and single-celled green algae (*Chlamydomonas*) for their ancestral characteristics. Our dataset’s combination of multiple species and diverse cell types gives a broad view of expressed proteins across *Viridiplantae*.

**Figure 1.**
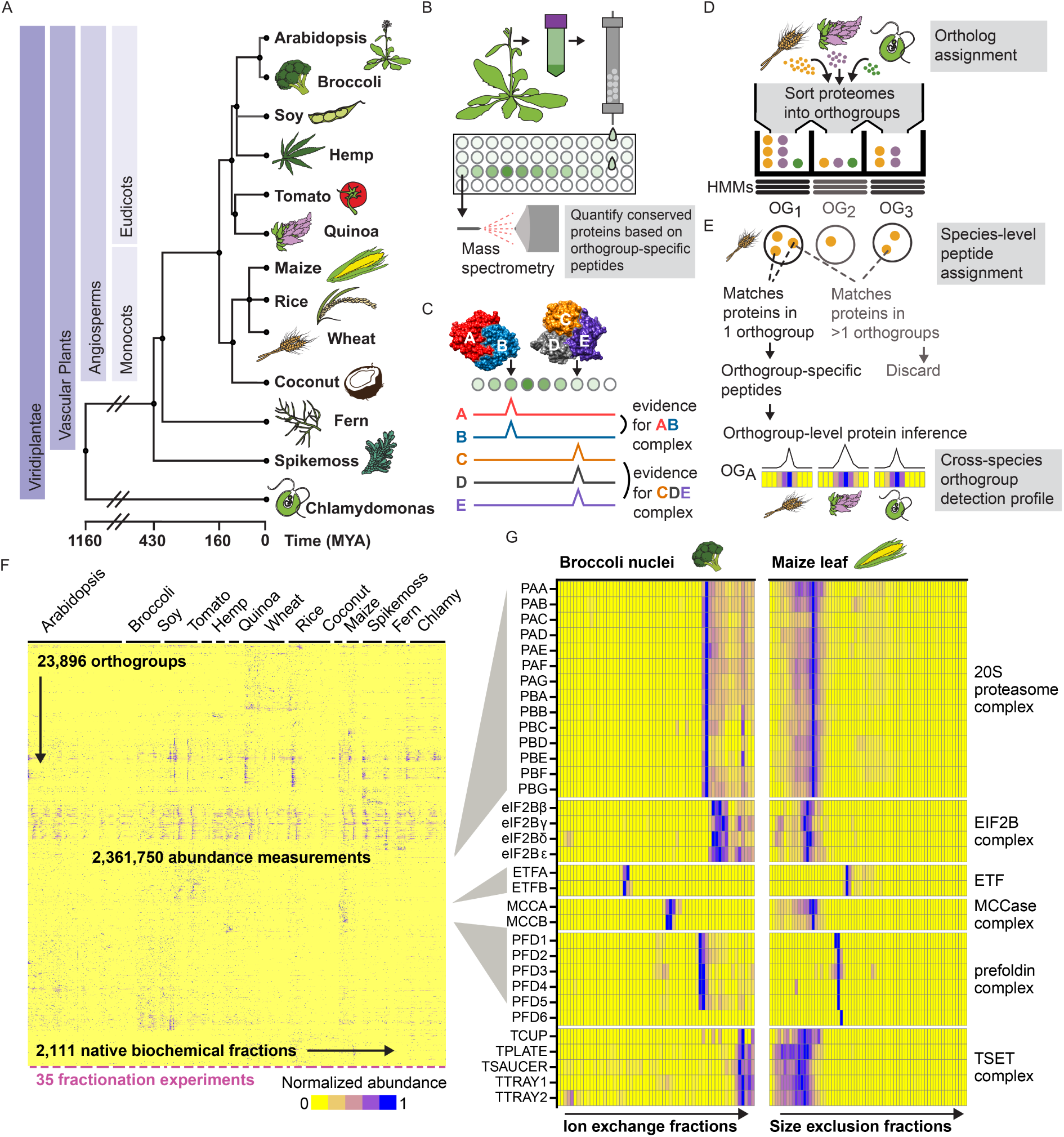
Integrative Co-fractionation Mass Spectrometry (CF-MS) Workflow Used to Determine Stable Plant Protein Complexes. See also Table S1. **(A)** The selected thirteen species represent a broad range of evolutionary time (MYA, million years ago (Kumar et al., 2017)). **(B)** Native extracts from each plant are chromatographically separated and the proteins in each fraction identified by mass spectrometry. **(C)** Co-fractionation of proteins is evidence of physical association. **(D)** Proteins from each species’ proteome are first assigned to orthologous groups (OGs). **(E)** Peptides that match more than one orthogroup (light gray text) are not used, however peptides that uniquely match a single orthogroup are used to quantify the abundance of an orthogroup in individual chromatographic fractions. Elution profiles for each orthogroup are graphically represented here as ridgelines or heat maps (blue showing abundance) across multiple separation experiments. **(F)** Heat map of the full dataset of abundance measurements for each of the 23,896 detected orthogroups across all fractionations for the thirteen species. Blue indicates non-zero signal. Dashes under heat map delineate each fractionation experiment. **(G)** Enlarged portions of **(F)** showing examples of strong co-elution observed for subunits (names at left, see Table S1) of six well-known protein complexes (names at right). Color intensity (blue is positive signal) depicts measured abundances for each orthogroup (labeled at left) in two distinct chromatographic separations (labeled at top) out of the 35 total separations.

We separated each native, non-denatured protein extract by some biophysical property, either size exclusion chromatography (SEC), ion exchange chromatography (IEX), or isoelectric focusing (IEF) (**Figure 1B-C**). Each chromatographic fraction was analyzed using high resolution, high sensitivity liquid chromatography/mass spectrometry (LC/MS). In all, we collected 14,520,970 interpretable peptide mass spectra from 2,111 individual chromatographic fractions, each capturing distinct subsets of native plant proteins and protein assemblies. These protein abundance measurements will be broadly valuable for addressing diverse questions of plant biochemistry and function, including protein modifications, expression, and determining which species or tissue contains high abundance of any particular protein.

### An evolution-informed strategy improves proteomics coverage and enables comparisons across species with different ploidy levels

Integrating protein observations from different organisms is complicated due to orthology mapping. This long-standing problem is even more extreme in plants due to their often complex and polyploid genomes, as well as past whole-genome duplications (Jiao et al., 2011). For example, most farmed wheat is hexaploid and contains over 100,000 genes, which complicates comparison to model diploids such as *Arabidopsis,* with its approximately 35,000 genes. The existence of multiple near-identical proteins also reduces the number of peptides that uniquely match a single protein, reducing protein recovery by standard proteomics methods. Current protein-grouping statistical methods for assigning peptides to proteins tend to perform erratically for highly redundant genomes, in practice allocating shared peptides semi-randomly across similar proteins. We thus developed a novel evolution-informed protein-grouping approach which is generally applicable to proteomic data from any arbitrary number of different species.

Our strategy was to interpret mass spectral observations in terms of protein orthogroups, rather than individual proteins. An orthogroup (OG) is a set of genes in modern organisms that all derive from the same original gene in those organisms’ last common ancestor. We began by assigning all protein-coding genes in each plant species to predetermined orthogroups (Huerta-Cepas et al., 2017), thereby organizing highly related protein sequences into homologous protein families (**Figure 1D**). We then considered mass spectra from any orthogroup member protein as evidence for the abundance of their orthogroup (**Figure 1E**). This analysis markedly increased the number of observed peptides because peptides shared by multiple members of an orthogroup (but not unique to a single protein) could now contribute to quantification. Importantly, orthogroup identifiers, unlike individual protein identifiers, are consistent across species and could be used as a key for integrating data from multiple species. Thus, we collapse the three proteins in the Ribosomal Protein L36 orthogroup in *Arabidopsi*s to a set, directly comparable to the behavior of that orthogroup in other species, for example the set of seven proteins in this orthogroup in wheat. (We refer to orthogroups by available *Arabidopsis* common gene names throughout; **Table S1** provides additional identifiers.) **Figure 1F** visually summarizes our over 2 million protein abundance measurements across plant species integrated into this comparative phylogenetic framework and **Figure 1G** highlights specific examples of complexes from the data in **Figure 1F** (described in **File S1,** data in **File S2**).

Consistent with ploidy levels (**Figure 2A**), diploid organisms such as *Arabidopsis* and *Chlamydomonas* exhibited a peak of one protein per orthogroup. In contrast, tetraploid quinoa and soy each showed a peak of two proteins per orthogroup (one from each of two subgenomes) and in the same manner, hexaploid wheat showed a peak of three proteins. Our data are consistent with the hypothesis that the current spikemoss (*Selaginella moellendorfii*) genome was a greenhouse hybrid (Banks et al., 2011), with a peak of two proteins per orthogroup, or one protein per parent genome.

**Figure 2.**
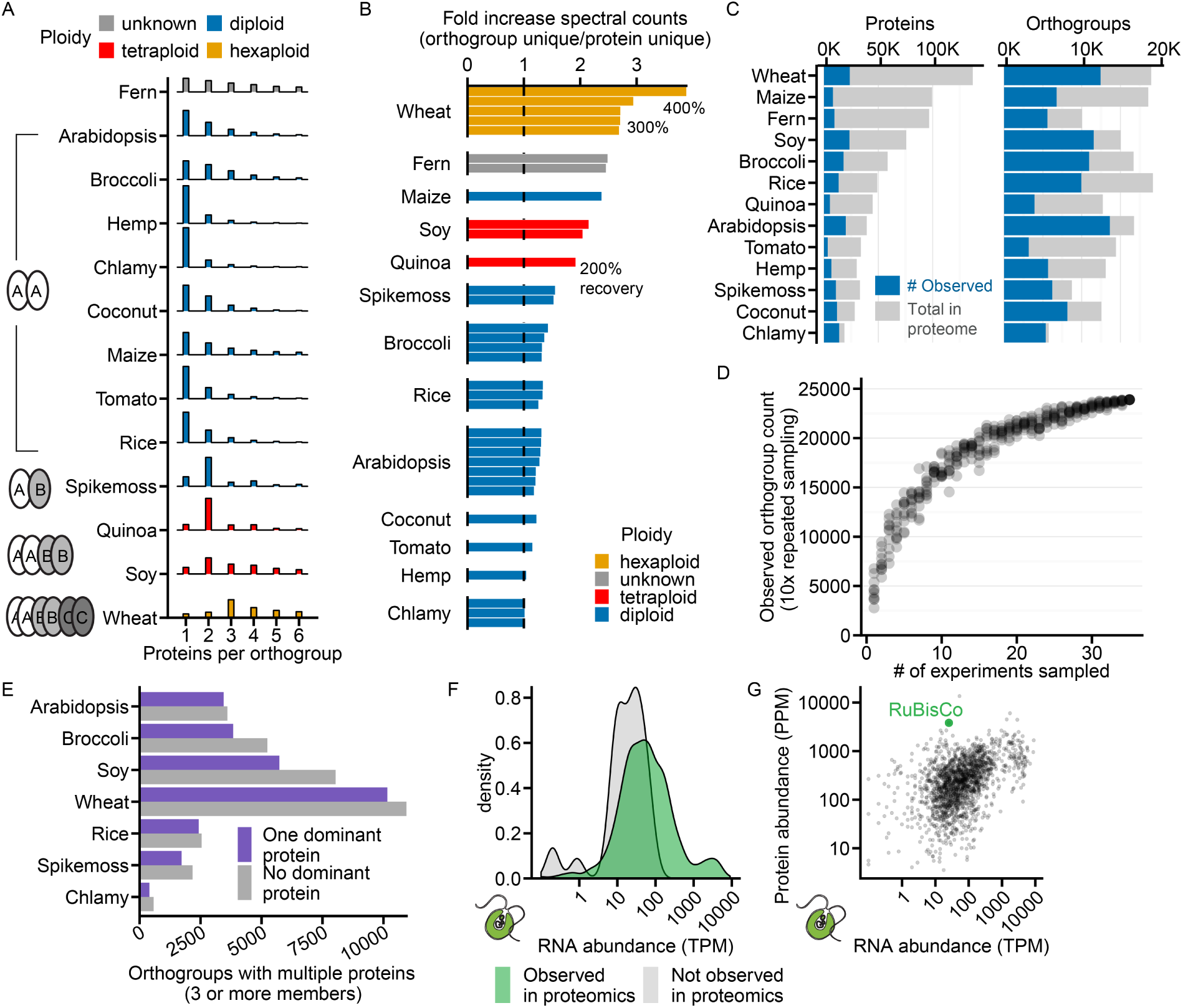
Proteomics in High-Ploidy Species Enhanced by Assignment of Proteins to Orthogroups. See also Figure S1. **(A)** Number of proteins assigned per orthogroup for each plant species in our study colored by ploidy. Shaded ovals at left represent subgenome organization. **(B)** Fold increase (x-axis) in peptide spectral matches that identify unique orthogroups vs. unique proteins. Each bar represents a single fractionation experiment conducted on the species named at left and color-coded by ploidy as in **(A)** **(C)** The number of observed proteins (left plot) or orthogroups (right plot) experimentally observed (in blue) compared to the possible total in the proteome (gray). Note that relative coverage per species is a function of the amount of data collected from that species in this study. **(D)** Our data set of 35 fractionations and 13 species is sufficient to identify the majority of orthogroups possible by this method. Each dot represents the number of orthogroups identified (y-axis) in a subsample of *n* experiments (x-axis), with sampling repeated ten times per each *n*. **(E)** Orthogroups with more than two proteins were approximately equally likely to be represented by a single dominant protein as not, regardless of ploidy. **(F)** Orthogroups observed by mass spectra (green) represent those with higher mRNA abundances (TPM, transcripts per million, log scale; data from (Panchy et al., 2014)), as shown for *Chlamydomonas*. Gray represents orthogroups not observed in our study. **(G)** Log-scale protein abundances (y-axis) show expected correlation with RNA abundances (x-axis, Transcripts per million; same as in **(F)**) in *Chlamydomonas*, however with numerous outliers, notably, RuBisCo (green dot).

This orthogroup-based proteomics interpretation strategy increased the recovery of unique spectral counts for the highly redundant proteome of hexaploid wheat by over 300%, while not strongly impacting organisms with small diploid genomes, e.g., *Chlamydomonas* (**Figure 2B**). Similarly, coverage of observed orthogroups is less variable than coverage of observed proteins across species. (**Figure 2C**). Collapsing sets of evolutionarily-related proteins into orthogroups is thus a flexible and broadly applicable solution for cross-species proteomics analyses, especially across plants with varying ploidy levels.

### Characteristics of expressed plant proteomes

Our data represent over 141,910 unique proteins and 23,896 orthogroups from diverse species and extracts, the largest proteomic survey of plants performed to date, covering broad areas of function (**Figure S1**). We sufficiently capture the proteome observable by native fractionation mass spectrometry, as evidenced by saturating the number of new orthogroups observed per additional experiment (**Figure 2D**), suggesting that more samples are unlikely to significantly improve conserved proteome coverage. In all, we observed 96.7% of the 11,339 most conserved *Viridiplantae* orthogroups (details in Methods). Our dataset, therefore, provides a meaningful snapshot of the conserved and expressed proteome of green plants.

Collapsing sets of proteins to orthogroups masks individual protein behavior, so we examined the behavior of paralogs within orthogroups. We found that about half of orthogroups with three or more member proteins contain one dominant protein that is highly expressed relative to other members (**Figure 2E**). An approximately equal set have protein expression more evenly shared across members. These trends were consistent across multiple species with widely varying genome sizes (**Figure 2E**). Thus, some degree of functional divergence, as measured by differential expression, was evident in about 50% of multi-gene orthogroups.

The proteins we observed also tended to be products of more abundant mRNAs (**Figure 2F**). Work in other organisms has shown that there is an imperfect correlation between protein abundances and RNA transcript levels, largely attributable to post-transcriptional, translational, and degradation rate effects on steady-state protein levels (Vogel and Marcotte, 2012). Our data here similarly illustrate this point (Spearman r = 0.45, **Figure 2G**). One notable outlier is the enzyme RuBisCo, the most abundant protein on earth (Feller et al., 2007) and the most abundant protein in 16/35 of our experiments. RuBisCo is substantially more abundant at the protein level than would be expected based on its transcript levels, e.g. in *Chlamydomonas* (**Figure 2G**) or in *Arabidopsis* tissues (**Figure S1**).

### Systematic identification and scoring of stable protein-protein interactions

In many cases, subunits of known complexes show co-elution patterns easily detected by eye, as for subunits of the 20S proteasome, prefoldin, and TSET complexes that respectively coelute with distinct, complex-specific, elution patterns (**Figure 1G**). However, a computational framework is necessary to identify co-eluting proteins systematically and at high-throughput.

To quantitatively score co-elution behavior indicative of stably interacting proteins, we employed a supervised machine learning approach based upon observed data for known complexes. Protein-protein interactions were derived solely from the coordinated separation behavior of proteins over multiple, orthogonal biochemical fractionation experiments. We required co-fractionation evidence in at least three species so as to capture well-supported and evolutionarily conserved interactions. We trained an Extra Trees classifier (with 5-fold cross-validation) to distinguish between characteristics of pairs of proteins known to stably interact and random pairs of observed proteins (details in Methods). The classifier assigned a probabilistic CF-MS score (Co-Fractionation-Mass Spectrometry score) between 0 and 1 to each potential protein-protein interaction (**File S3**), with 1 indicating a high likelihood of physical association based on observing strongly coordinated protein elution profiles, and 0 indicating no evidence for interaction.

We next aimed to rigorously assess statistical confidence for physical protein-protein interactions, evaluating against a fully withheld test set of 886 known protein interactions that were not used at any point during model training. Ranking protein interactions by their CF-MS scores accurately recapitulated this withheld test set (**Figure 3A**) and allowed us to measure classifier error rates: For interactions with CF-MS scores over 0.50, we observed 90% precision (*i.e.* ≤10% false positive interactions) and 23% recall. Interactions with a CF-MS score above 0.2 exhibited 50% precision at 51% recall, and thus were still informative in many cases (**Figure 3B**).

**Figure 3.**
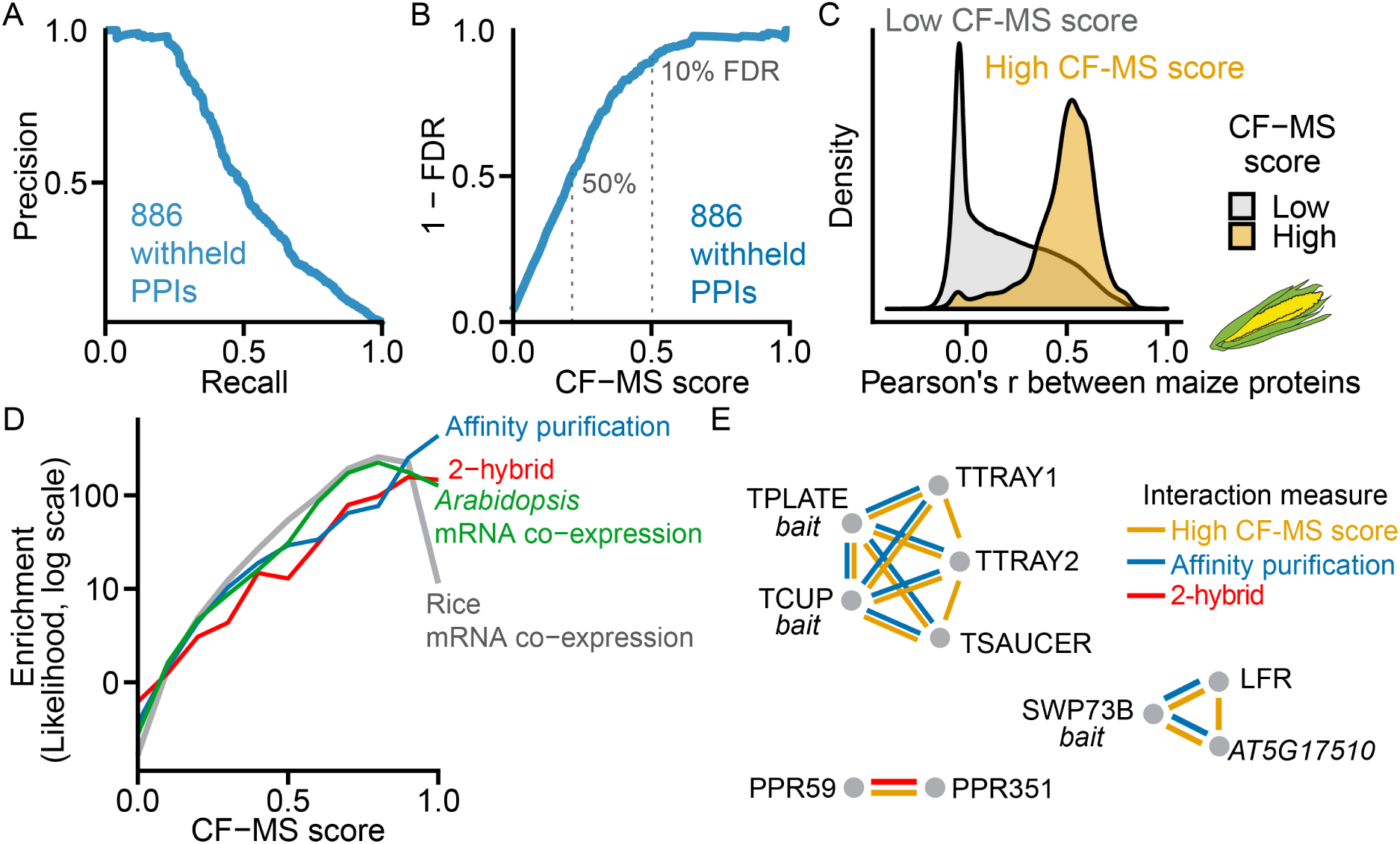
Derivation and Global Validation of Protein Co-Complex Interactions. **(A)** Precision-Recall of CF-MS scored protein-protein interactions (PPIs) on 886 known interactions withheld from training. **(B)** False Discovery Rate (FDR) vs. CF-MS scores for the same withheld set as in **(A)**. **(C)** PPIs with high CF-MS scores (FDR < 10%) are highly correlated in a species withheld from training (maize). **(D)** Protein interactions with higher CF-MS scores were more likely to have been identified by affinity purification, 2-hybrid in *Arabidopsis*, and were more likely to be co-expressed in *Arabidopsis* and rice. **(E)** Agreement of the CF-MS protein-protein interactions (yellow) with affinity purification (blue) and 2-hybrid interactions (red) for three protein complexes.

### Confirming CF-MS interactions by independent assays and chemical cross-linking

Our measured high-confidence protein interactions agree with independent protein interaction observations. As a direct test of our method, we asked if independent CF-MS data from maize, a plant species that was not used to train the model, showed the same trends. Ortholog pairs with a high CF-MS score (>0.5, derived from non-maize plants) co-eluted strongly in a fractionation experiment of maize shoots (**Figure 3C**). Thus, CF-MS-supported interactions are highly concordant with biochemical separation behavior of proteins in an unassayed plant species. We also compared our observed interactions to other independent plant protein interaction datasets, finding that protein pairs with higher CF-MS scores were more likely to affinity purify together, to interact by yeast-2-hybrid, and to exhibit coordinated mRNA expression both in *Arabidopsis* and in rice (**Figure 3D)**. Our high confidence scores align with and provide orthogonal support for biologically interesting interactions previously discovered by AP-MS and yeast-2-hybrid assays. **Figure 3E** illustrates three such cases for the TPLATE complex proteins (Gadeyne et al., 2014), the interactions between SWI/SNF components with LEAF AND FLOWER RELATED (LFR) and an uncharacterized protein AT5G17510 (Vercruyssen et al., 2014), and the interaction between two uncharacterized pentatricopeptide repeat proteins PPR59 and PPR351 (Arabidopsis Interactome Mapping, 2011).

To further independently validate our derived protein complexes using unbiased orthogonal methods (and thus avoid “cherry-picking” promising test cases), we implemented two untargeted, large-scale biochemical approaches. For the first method, we plotted expected monomeric mass against observed mass in a representative *Arabidopsis* size exclusion experiment and demonstrated that a substantial proportion of proteins eluted with a mass considerably larger than their monomeric masses, supporting that endogenous complexes remained intact in our experimental conditions (**Figure 4A**, Methods).

**Figure 4.**
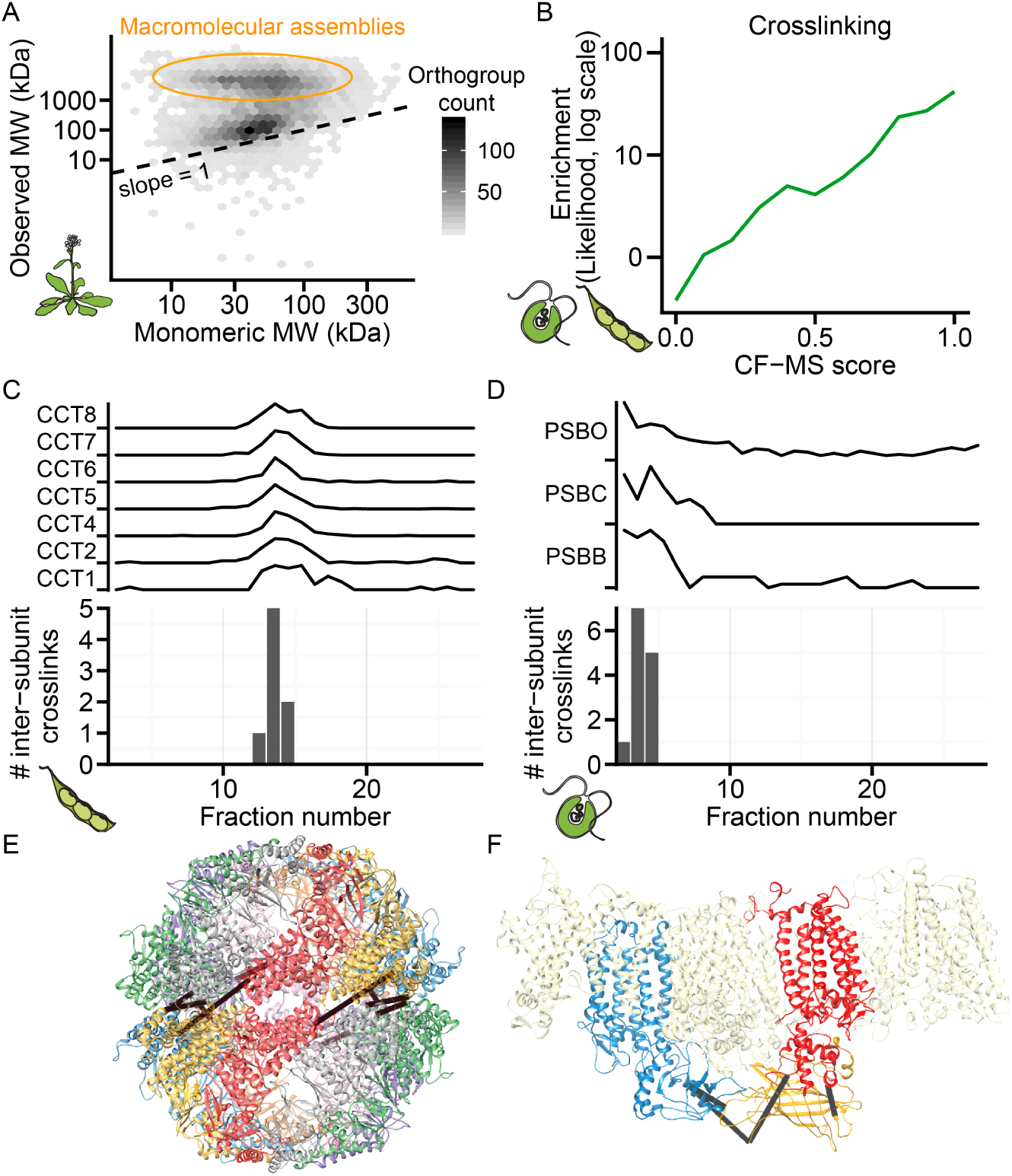
Protein Complexes Validated by Calibrated Molecular Mass Determination and Direct Chemical Cross-linking. **(A)** Observed mass vs. predicted monomeric mass in a representative *Arabidopsis* size exclusion chromatography (SEC) fractionation. Shading reflects the number of orthogroups per hexagonal bin. **(B)** Cross-linked proteins from soy and *Chlamydomonas* are more likely (green line, log-likelihood) to have high CF-MS scores compared to non-cross-linked observed proteins. **(C-D)** Inter-subunit cross-links only appear in fractions where complex subunits co-elute. Elution profile and inter-subunit crosslinks for soy T-complex chaperonin (CCT) shown in **(C)** and *Chlamydomonas* Photosystem II (PSB) shown in **(D)** **(E-F)** 3D homology models of complexes (see Methods) with observed inter-subunit cross-links (black lines). Soy CCT **(E)** is colored by subunit, and *Chlamydomonas* Photosystem II **(E)** highlights PSBB, PSBC, and PSBO as blue, red, and yellow, respectively.

For the second validation method, we performed global chemical cross-linking on fractionated protein extracts from soy sprout and *Chlamydomonas*, covalently tethering pairs of interacting proteins within each chromatographic fraction. We identified 194 heteromeric protein-protein interactions from soy and 228 from *Chlamydomonas*, at a false discovery rate of 1% and ΔXlinkX score ≥70 (**File S4**). Cross-linking recovered 31 withheld test set positive interactions and one negative interaction, empirically estimating the false discovery rate at 3%. We observed high concordance between the cross-linked protein pairs and our CF-MS determined interacting proteins, and protein pairs with high CF-MS scores were considerably more likely to be cross-linked (**Figure 4B**).

Furthermore, our observed cross-links were consistent with physical constraints of known protein interactions (**Figure 4C-F**). For example, intersubunit cross-links in the soy CCT chaperonin complex were exclusively identified in the same fractions as co-eluting CCT subunits (**Figure 4C**) and occurred between residues appropriate to the cross-linker length (<30Å Cα-Cα) (**Figure 4E**). While this complex is well documented in animals, a 22S complex containing CCT subunits was only reported in 2019 in *Arabidopsis* (Ahn et al., 2019). Our data confirm that the CCT complex is conserved across plants and indicate that the specific 3D subunit organization in plants resembles that in animals. Similarly, co-fractionation and cross-linking of the Photosystem II complex was observed in *Chlamydomonas*, with the observed inter-protein cross-links located at appropriate adjacent solvent-accessible subunit interfaces (**Figure 4D, F**) as expected for the native conformation of this complex. The recovery of multiple structurally coherent inter-subunit cross-links between co-fractionating subunits provides experimental evidence that CF-MS faithfully captured protein assemblies.

Thus, a combination of direct experimental validation using independent biochemical methods (co-fractionation in an independent organism, calibrated size exclusion chromatography, and chemical cross-linking) and comparison to independently determined protein interactions from the literature all indicate that the CF-MS measurements provide strong evidence for physical assemblies.

### Identification of multiprotein complexes confirms those inferred by gene content and reveals additional novel assemblies

As our CF-MS datasets faithfully captured many large multiprotein assemblies (**Figures 1G, 4A, C-F**), we next sought to systematically define higher-order plant protein complexes by clustering the proteins based on the measured pairwise interactions at FDR < 10% (**Figure 5**). Instead of choosing a single clustering cutoff to define discrete complexes, we selected multiple cutoffs to reflect hierarchies of interacting proteins. For example, one cutoff defined the 80S ribosome, while a finer cutoff differentiated its 40S and 60S subcomplexes (**Figure 5**). Orthogroups in well-characterized complexes are annotated dark green; however, we identified several previously unreported subunits and interactors with the potential to enrich our understanding of how these known complexes function in plants. Excitingly, we observed many complexes comprised of novel associations (**Figure 5**, yellow) as well as proteins that remain uncharacterized in plants (**Figure 5**, bold circles). Our clustered and annotated set of high confidence protein complexes is available as **File S5.**

**Figure 5.**
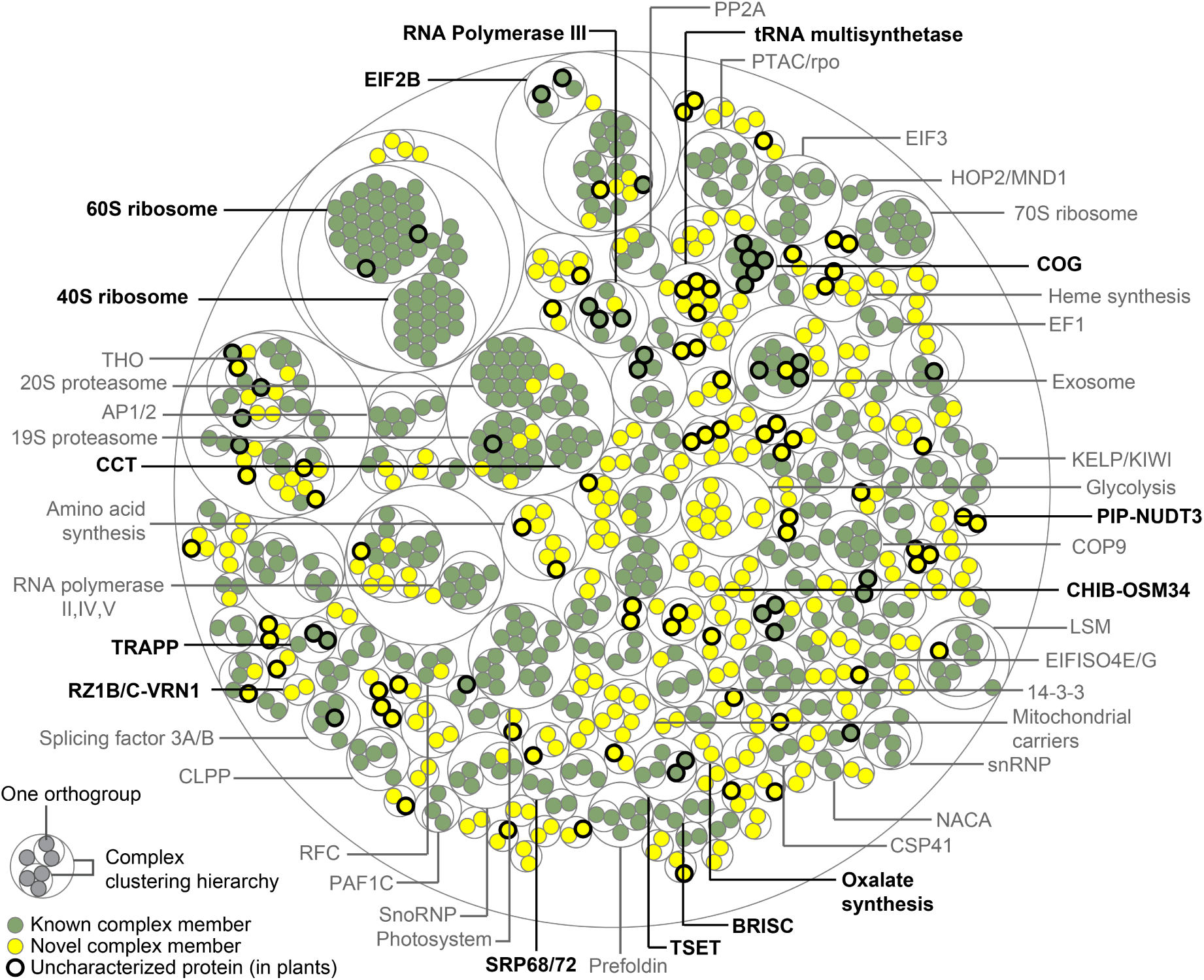
Overview of Evolutionarily Conserved Plant Protein Complexes. Thin concentric circles show the clustering hierarchy of protein-protein interactions into complexes for each of four clustering thresholds (see **File S5** for complex membership and annotations). Protein orthogroups (filled circles) are colored green for associations previously reported in any species or yellow for those first reported in this study. Bold outlines denote proteins uncharacterized in plants, defined as uncharacterized if all proteins in the orthogroup lack an *Arabidopsis* gene symbol and a Uniprot Function annotation. Bold complex labels are discussed in the text.

As internal positive controls, we identified 117 complexes that are reported in the literature. Some of these eukaryotic complexes, such as the Conserved Oligomeric Golgi (COG) complex, the SRP68/72 heterodimer, and the TRAPP and BRISC complexes have to our knowledge only previously been inferred in plants by gene content. Similarly, we find orthologs of complex members that have only been reported in non-plant species, such as a MAA3 (a plant ortholog of the yeast protein Sen1 and human Senataxin) interacting with RNA polymerase III, an interaction recently found in yeast to regulate RNA polymerase III termination (Rivosecchi et al., 2019).

Likewise, while homologs to the yeast and mammalian oligosaccharyltransferase (OST) complex subunits exist in plants, the full complex has not been biochemically isolated (Strasser, 2016). We observe a plant OST complex with overlapping membership to the yeast and mammalian OST complex, and detect cross-links between HAP6 and OST48 in soy, suggesting the plant OST complex resembles that of other eukaryotes. We identify potential novel OST components Stomatin-like protein 1 (SLP1), SPC25, and EMC1. We also identified components of the initiation factor (eIF)2B complex in which the eIF2Bγ/ε and eIF2Bβ/δ dimers were clearly co-purifying with each other (**Figure 1G**), but the eIF2Bα subunit appeared to be less stably associated. The existence of the eIF2B complex in plants has been speculative as this complex has not been characterized and isolated from plants, in contrast to its well-established observation and characterization in other eukaryotes (Browning and Bailey-Serres, 2015).

Protein interactions provide a means to characterize, corroborate, and predict protein functions *via* guilt-by-association. We found several instances where a top-scoring interaction to an uncharacterized protein could be confirmed in the literature, serving as additional positive controls. For example, the most confident interactor with the recently characterized *Arabidopsis* protein AT5G14910 is the chloroplast ribosome protein RPS1; the interaction was independently confirmed in a recent study (Pulido et al., 2018). We also observed instances of interactions between proteins catalyzing consecutive enzymatic reactions, such as a novel interaction between OXP1 and GEP, enzymes catalyzing the last two steps of glutathione degradation. In cases where complexes identified at a stringent FDR lack known members, using the core subunits to query scored interactions often recovers the expected subunits and potential novel interactors. Because protein interactions can lead to functional insights, we provide a public tool for the community to query the CF-MS interactions at http://plants.proteincomplexes.org.

### Alternative multiprotein assemblies are apparent in the plant lineage

Analysis of protein complexes has revealed that homologous gene products are not always assembled in the same way (Marsh and Teichmann, 2015). We found many cases in which plants appear to have alternate arrangements of interacting proteins relative to other lineages, including the adoption of plant-specific subunits. Furthermore, we identified cases in which plants exhibited analogous assemblies to those in other lineages, achieving similar architectures or functions, but through distinct molecular interactions. In both cases, protein sequence homology alone was inadequate to predict protein complex structure or function without knowledge of the assemblies themselves.

One prominent example is a conserved tRNA multi-synthetase complex (MSC). While functional assemblies of essential aminoacyl tRNA synthetases seem ubiquitous across all life forms, distinct architectures, memberships, and accessory proteins have been cataloged in diverse organisms including animals, yeast, archaea, and bacteria (Laporte et al., 2014). They are frequently loosely associated and condition-dependent, making complete composition difficult to define. A candidate MSC was recently reported in *Arabidopsis* (McBride et al., 2019). We observed a conserved megadalton-scale MSC with architecture and accessory subunits distinct from those in animals, fungi, and bacteria but with notable parallels (**Figure 6A**). Our plant MSC lacks the p38, p43, and p18 scaffolding subunits critical for assembly of the human multi-synthetase. Instead, it contains the plant ortholog of ARC1, the central organizing cofactor of the yeast multi-synthetase. Conservatively, the plant MSC complex appears to contain the ortholog of ARC1, Ybak (a member of the trypanosome MSC (Cestari et al., 2013)), Clustered Mitochondria Homolog (CLU), the WD40 scaffold protein VIP3/SKI8, and a distinct set of tRNA synthetases, among them glutamate, isoleucine, and tryptophan-tRNA ligases (E, I, and W). Peripheral members may include valine, tyrosine, histidine, aspartate, proline, threonine, leucine, glutamine, and methionine-tRNA ligases (V, Y, H, D, P, T, L, Q, and M) (**Figure 6A**). Of the 20 eukaryotic tRNA ligases, 8/9 observed in the mammalian MSC (all but asparagine) appear to be in the plant MSC complex, suggesting that there could be an advantage to assembling these particular tRNA-ligases.

**Figure 6.**
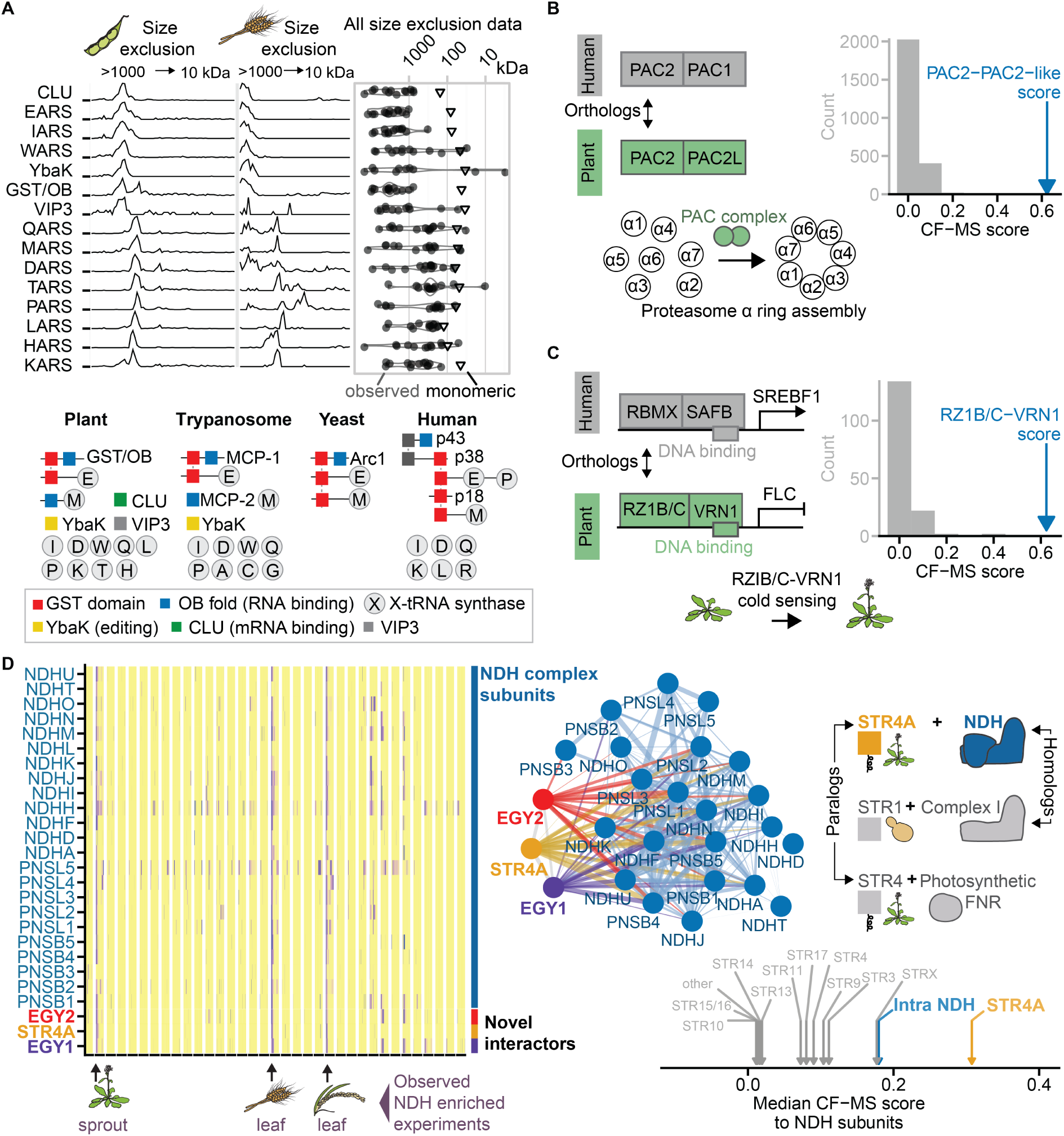
Alternative Assemblies in Plant Analogs of Animal Multi-Protein Complexes. **(A)** Plant Multi-tRNA Synthetase Complex (MSC). Top left, elution profiles of proteins observed in large molecular weight complexes containing aminoacyl tRNA synthetases in soy and wheat size exclusion fractionations. Top right, the observed molecular mass (circles) for each protein at left in all plant size exclusion separations in our data set compared to the predicted monomeric mass (triangles). Bottom, schematic of the domain structure and organization of MSC proteins in representative eukaryotic lineages. **(B)** A plant proteasome assembly chaperone complex, with orthology of plant PAC2 to the human analog PAC2 indicated with double-headed black arrow. Right, the CF-MS score for the PAC2-PAC2L interaction (blue arrow) far exceeds that of any other protein interaction score with either PAC2 or PAC2L. Gray bars are binned CF-MS interaction scores for all other protein interactions. **(C)** A plant transcriptional response module, with orthology of RZ1B/C to the human analog RBMX indicated with double-headed black arrow. Right, the CF-MS score for the RZ1B/C-VRN1 interaction (blue arrow), gray bars as in **(B)**. **(D)** Novel subunits of chloroplast NADH dehydrogenase-like complex (NDH). Left, heatmap shows coelution (purple) of known NDH subunits along with three novel interactors in specific plant extracts (arrows below). Middle, network diagram with proteins (circles) connected by interaction lines where line thickness reflects CF-MS score. Right, illustration of conserved molecular architecture and use of rhodanese sulfurtransferase subunit modules in electron transport complexes - two plant-specific (NDH, FNR) and one conserved mitochondrial (Complex I). Bottom, median CF-MS scores to all NDH subunits shown for known NDH subunits and all rhodanese-like domain proteins in plants.

Just as the presence of orthologs could not predict the plant MSC complex, the genetic absence of orthologs does not predict the absence of functionally similar complexes. One example is the complex of proteasome assembly chaperones. In humans, a stable heterodimer of PAC1 and PAC2 aids the assembly of the proteasome alpha subunit ring (Hirano et al., 2005). While plants lack a PAC1 gene, we found a plant-specific PAC2-like protein associated with PAC2 (**Figure 6B**). No proteasome assembly chaperone complex has been previously described in plants, suggesting that the PAC2/PAC2-like complex likely performs this function.

We also found several instances where plants utilize lineage-specific subunits to co-opt known molecular modules to serve plant-specific functions. One example is a heterodimer of a conserved eukaryotic transcription factor. In humans, RBMX interacts with SAFB to bind the promoter of the SREBP1 gene to regulate sterols in the liver (Omura et al., 2009). We found the plant ortholog of the RBMX transcription factor (RZ1B/C) interacting with the plant-specific protein VERNALIZATION1 (VRN1) (**Figure 6C**), both of which are known to regulate the *FLOWERING LOCUS C* (*FLC*) gene (Levy et al., 2002; Wu et al., 2016) and together control a crucial plant-specific event: rapid flowering at the appropriate seasonal time.

Finally, the chloroplast NADH dehydrogenase-like (NDH) complex provides a more intricate example of how plants have adapted conserved architectural modules with plant-specific proteins to serve plant-specific purposes. NDH is a chloroplast complex that shares architecture with the respiratory Complex I of mitochondria, with both complexes functioning in electron flow (Shikanai, 2016). We identified known NDH subunits and found three additional subunits, EGY1, EGY2, and STR4A (**Figure 6D, left**). These new subunits have high CF-MS scores for interactions with multiple subunits of the NDH complex, scoring highly with NDH subcomplex B and L members, and in some cases scoring higher than the interactions between known members of the NDH complex itself (**Figure 6D, network**). EGY1 and EGY2 are chloroplast-localized intramembrane metalloproteases, but their specific functions remain unknown (Adamiec et al., 2017). The specific enrichment of EGY1, EGY2, and the NDH complex in Bundle Sheath chloroplasts of C4 plants supports the association of these metalloproteases with NDH *in vivo* (Majeran et al., 2008). The third new subunit, STR4A, is a rhodanese-like protein of unknown function. While mitochondrial Complex I is known to interact with an accessory rhodanese domain sulfurtransferase, no such subunit has yet been reported for the architecturally similar plant NDH complex. STR4A is one of 6 rhodanese-like domain proteins in *Arabidopsis*; however, the CF-MS scores indicate that the association of STR4A with the NDH complex is unique among those six (**Figure 6D, bottom right**). One clue to the role of STR4A may come from the related protein, STR4, required for localization of the photosynthetic FNR complex to the thylakoid membrane of chloroplasts (Lintala et al., 2014). However, because NDH, FNR, and Complex I are all electron transport complexes, it is also possible that sulfurtransferase activity is important for some shared function in electron transport.

These observations highlight how shared features such as orthology and protein complex architecture can be informative, but as plants have alternatively purposed many proteins and complexes, directly measuring specific protein interactions and assemblies is necessary to understand the functional roles of plant proteins.

### Interaction-to-phenotype: Discovering protein functions and phenotypes from protein interactions

Ultimately, the utility of large-scale datasets is their ability to drive biological discovery. Interacting proteins are more likely to share phenotypes (Fraser and Plotkin, 2007; Lage et al., 2007; McGary et al., 2007) (**Table S3**) and thus our data should provide a basis for linking genotype to phenotype and gaining biochemical insights into shared phenotypes. We therefore interrogated our dataset for cases where the derived interactions might suggest new hypotheses regarding protein function and phenotypes.

We found several links between pathogen defense and immunity genes through protein interactions. Plant pathogen resistance mechanisms are an area of intense research, as pathogens cause billions of dollars in crop losses annually (Savary et al., 2019). We discovered two novel complexes related to plant-pathogen interactions. The first is comprised of Basic endochitinase B (CHIB) and Osmotin-like protein 34 (OSM34) (**Figure 7A**), representing two different protein families, Pathogenesis-Related protein group 3 (PR3) and Pathogenesis-Related protein group 5 (PR5) respectively. Each protein has been individually reported to target fungal cell walls, and both are highly co-expressed following infection by the gray mold fungus *B. cinerea* (Dhawan et al., 2009). To our knowledge, this is the first report of a stable protein complex among members of different pathogenesis-related protein classes. Mechanistic characterization of this protein complex could aid the development of strategies to prevent devastating crop losses from fungal infection.

**Figure 7.**
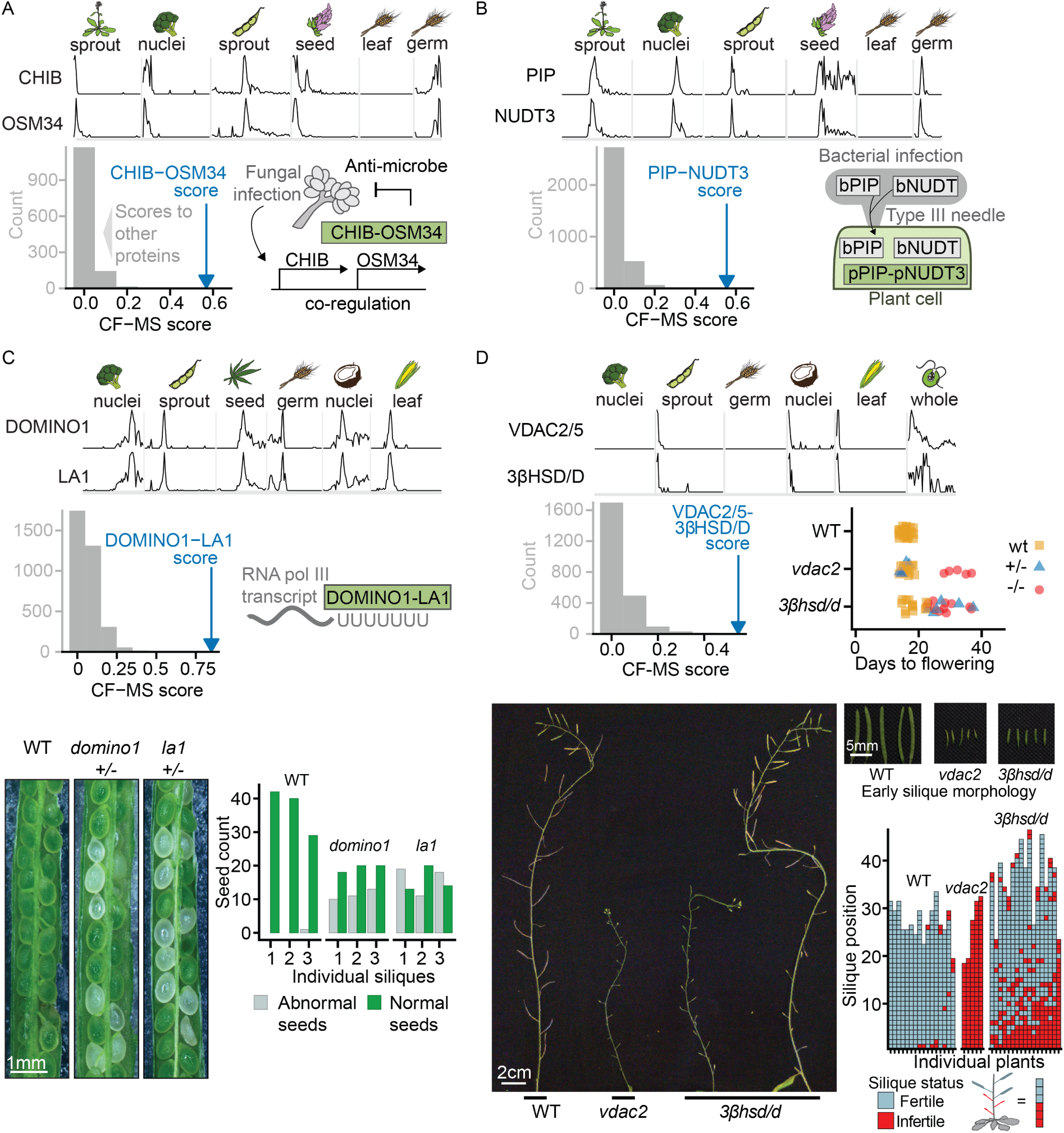
Connecting Plant Genes to Phenotypes *via* Their Interactions See also Table S3. The top section of each lettered panel shows sparklines with sample species and tissues indicated above. Bottom left panel of each lettered panel shows the complex interaction score (CF-MS) between subunits (blue arrow) is far greater than the interaction of either subunit with any other observed protein (gray bars representing binned scores). **(A)** OSM34 and CHIB form a complex, consistent with co-expression evidence in response to fungal infection (diagram bottom right). **(B)** PIP and NUDT3 form a complex in plants. Bacterial members of the PIP and NUDT families are injected into plant cells by a Type III secretion system to modulate the plant immune response. **(C)** DOMINO1 and LA1 form a plant-specific ribosomal RNA-binding complex and heterozygotes of each have a similar T-DNA insertion mutant phenotype of abnormal white seeds containing arrested embryos. Bottom left, representative portions of siliques from genotypes as labeled. Lower right, quantification of visually abnormal seeds in three siliques of each genotype. Ratio of normal to abnormal seeds reflects variable penetrance of the mutant phenotype and presence of homozygous and heterozygous embryos in each silique. **(D)** Arabidopsis plants homozygous for VDAC2/5 or 3βHSD/D T-DNA insertion mutants show delayed flowering and reduced number of fertile siliques compared to wild type plants of the same stage. Lower panels illustrate fertility defects with main inflorescences at end of flowering. While *vdac2* homozygotes produce almost no seeds, *3βhsd/d* mutants show a range of fertility levels, ranging from plants with almost no seed-containing siliques to plants in which only early siliques show fertility defects. An enlarged view of wild type and early infertile siliques from plants of the genotypes is shown.

A second novel pathogen-related protein complex contains proline iminopeptidase (PIP) and Nudix hydrolase 3 (NUDT3) (**Figure 7B**). The native role of these proteins in plants is unclear, but intriguingly, bacterial versions of both PIP (Kan et al., 2018) and Nudix hydrolase (Dong and Wang, 2016; Mukaihara et al., 2004) are injected by bacterial type III secretion systems into plant cells in order to suppress plant immunity. While work remains to determine the native role of plant PIP and NUDT3, the information that these proteins form a stable endogenous complex will guide future efforts. The direct observation of known pathogen resistance proteins in complexes creates a concrete framework for interpreting previous results and examining the mechanism by which they impact plant health.

We also considered a new interaction between the DOMINO1 and LA1 proteins which served to confirm a joint role for the interaction partners in embryonic development. Loss of function of either of these genes is known to produce a nucleolar hypertrophy phenotype and nonviable embryos (Fleurdépine et al., 2007; Lahmy et al., 2004). While LA1 is an RNA-binding protein found in diverse eukaryotes, DOMINO1 is a plant-specific protein with mutations linked to ribosome biogenesis defects and slow embryonic growth (Lahmy et al., 2004). We observed a stable DOMINO1/LA1 complex in multiple plants and tissues (**Figure 7C**) and confirmed that both subunits affect the same biological process by comparing the phenotypes of individual *domino1* or *la1* insertional mutant lines of *Arabidopsis*. Heterozygous *domino1* or *la1* mutant plant lines produced siliques with many abnormal clear embryos (**Figure 7C**). These phenotypic similarities support a functional complex of the DOMINO1 and LA1 proteins *in vivo* with a proposed role in ribosome biogenesis.

Protein interactions also provide a basis to predict entirely new phenotypes. We exploited this trend for a new interaction between the mitochondrial outer membrane porin VDAC2/5 and 3β-hydroxysteroid dehydrogenase/C-4 decarboxylase (3βHSD/D). As *vdac2* mutants have a well-characterized late flowering and sterility phenotype (Tateda et al., 2011), we predicted by interaction that *3βhsd/d* mutants would show similar defects. We directly compared the phenotypes of *Arabidopsis* bearing loss of function T-DNA insertions in VDAC2 or 3βHSD/D2. Disruption of either gene delayed flowering, induced wavy leaves, and reduced fertility. (**Figure 7D**). The underlying defect of these shared phenotypes is likely related to transport and modification of plant sterols, as 3βHSD/D is a sterol modifying enzyme (Kriechbaumer et al., 2018), and VDAC2 is essential in mice for steroidogenesis (Prasad et al., 2015).

## CONCLUSIONS

By using mass spectrometry proteomics to define the major protein complexes shared across plants, we have constructed a reference map of the basic biochemical ‘wiring diagram’ of a plant cell. The data capture over two million protein abundance measurements from multiple tissues and diverse species, revealing stable protein complexes conserved over more than a billion years of plant evolution. The resulting map thus provides a global snapshot of protein organization in plants. While we have presented examples of paths to connect gene products with phenotypes and to test specific functional hypotheses, this rich dataset can be mined in myriad additional ways, laying the foundations for extensive basic and applied research across the vast landscape of plant biology.

## ACKNOWLEDGMENTS

The authors gratefully acknowledge Angel Syrett for plant illustrations, Benjamin Liebeskind for assistance with fern transcriptomics and feature calculation, Anna Battenhouse for data deposition assistance, Claire Palmer for early orthology calculations, Hong Qiao, Alan Lloyd, and the University of Texas UTEX Culture Collection of Algae for providing plant and algal samples, Dan Boutz for mass spectrometry assistance, Riddhiman Garge and Momo Wisath Sae-Lee for assistance with coconuts, Andrew Emili for early discussions and assistance, and John Wallingford and Dannie Durand for constructive feedback and editing. The research was funded by grants from the Welch Foundation (F-1515 to E.M.M.), National Science Foundation (1237975 to P.C.R. and E.M.M.), Army Research Office (W911NF-12-1-0390), and National Institutes of Health (GM123683 to C.D.M., K99 HD092613 to K.D., R01 GM109076 to Z. J. C., and R35 GM122480 to E. M. M.).

## AUTHOR CONTRIBUTIONS

Conceptualization and Methodology, CDM, OP, EMM; Software, CDM, KD; Investigation, OP, CDM, CW, RMC, VJ, OXD; Formal Analysis and visualization, CDM; Writing – Original Draft, OP, CDM, EMM; Writing – Review & Editing, CDM, OP, VJ, OXD, KSB, ZJC, PCR, EMM; Funding Acquisition, CDM, KD, ZJC, PCR, EMM; Fern transcriptome, CDM, TK, MLS, SJR, EMM; Resources, SJR, ZJC, KSB, PCR, EMM; Supervision, EMM

## METHODS

### Data and code availability statement

The interaction data are publicly accessible *via* a dedicated web portal (http://plants.proteincomplexes.org). All raw and interpreted mass spectrometry data were deposited to the ProteomeXchange http://www.proteomexchange.org/ *via* the PRIDE (Perez-Riverol et al., 2019) partner repository with identifiers (PXD012810, PXD012865, PXD012969, PXD013004, PXD013080, PXD013093, PXD013198, PXD013213, PXD013214, PXD013264, PXD013280, PXD013281, PXD013282, PXD013300, PXD013320, PXD013321, PXD013322, PXD013369 PXD013704, PXD013735, PXD014617). Full documentation of computational analyses, non-external analysis scripts, and project data files are deposited at Zenodo (DOI:10.5281/zenodo.3466034), and analyses made use of the following github.com/marcottelab repositories: MS_grouped_lookup, protein_complex_maps, run_TPOT, MSblender, and run_MSblender. Fern (*Ceratopteris richardii*) transcriptome data and assemblies were deposited into the European Nucleotide Archive http://www.ebi.ac.uk/ena/ with accession number PRJEB33372, and proteome deposited at Zenodo (DOI:10.5281/zenodo.3467770).

### Native protein extraction

#### Tissues used

Broccoli heads (*Brassica oleracea var. italica*) and quinoa (*Chenopodium quinoa*) seed (husk removed) were purchased from Central Market H-E-B (Austin, TX), and hemp hearts “Raw Shelled Hemp Seed” (*Cannabis sativa*) from Trader Joe’s (Austin, TX). Leaves were harvested from young broccoli and tomato (*Solanum lycopersicum*) plants purchased at Barton Springs Nursery (Austin, TX) and from 8-day old (V2 stage) maize seedlings (*Zea mays*). *Selaginella moellendorfii* was purchased from Plant Delights Nursery, Inc., (Raleigh, NC). The following samples were provided by colleagues at the University of Texas at Austin: *Arabidopsis thaliana* Col-0 seedlings grown 5 days with a cycle of 16 hours light and 8 hours dark, or 3 days in the dark (Dr. Hong Qiao), *Arabidopsis thaliana ttg1-1* mucilage-less mutant seeds ((Koornneef, 1981), provided by Dr. Alan Lloyd), maize sprouts (*Zea mays)* (Dr. Jeffrey Chen), wheat (*Triticum aestivum)* germ extract, wheat seedlings, and *Arabidopsis thaliana* sprouts (Dr. Karen Browning), fern (*Ceratopteris richardii, Hn-n)* fronds, soy (*Glycine max)* seven day old sprouts (Dr. Stan Roux), and the algae *Chlamydomonas reinhardtii* UTEX90 (UTEX Culture Collection of Algae). Rice leaves (*Oryza sativa ssp japonica,* Kitaake cultivar) were provided by Dr. Pamela Ronald (UC Davis).

Unless otherwise stated, samples were quick-frozen and ground to a fine powder in liquid nitrogen using a chilled mortar and pestle. Powder was thawed to 4°C, and resuspended in approximately an equal volume of the specified lysis buffer plus protease and phosphatase inhibitors followed by nutation 30 minutes at 4°C. All subsequent steps were at 4°C. The crude homogenate was clarified by a low-speed spin (~3000 × g, 10 minutes), and the supernatant further clarified by a high-speed spin (~14,000 × g, 10 minutes). Protein concentration was determined by Bio-Rad Protein Assay or Bio-Rad DC Protein Assay, and 1-4 mg of extract was 0.45µ filtered (Ultrafree-MC HV Durapore PVDF, Millipore) and fractionated by HPLC chromatography. In general, detergent and salt were minimized where possible to avoid perturbing protein assemblies. Because the highly abundant plant protein RuBisCo dominates green tissues, we included etiolated seedlings (depleted for photosynthetic proteins) and two non-green tissues: germ tissues (enriched in proteins related to core cell biology), and isolated nuclei (enriched for DNA/transcription-related proteins) (**Figure S1**).

#### Green plant tissue extracts

Liquid nitrogen powders of leaves and/or sprouts of *Arabidopsis*, rice, wheat, broccoli, soy, tomato, fern, and *Selaginella moellendorfii* were lysed in Plant Lysis Buffer: 50 mM Tris-HCl pH 7.5 at room temperature, 150 mM NaCl, 5 mM EGTA, 10% glycerol, 1% NP40, 1 mM DTT, plant-specific protease inhibitors (Sigma) and phosphatase inhibitors (PhosSTOP EASY, Roche).

#### Embryonic plant tissue/seed extracts

Liquid nitrogen powders of quinoa, hemp seed, or *Arabidopsis thaliana* mucilage-less mutant *ttg1-1* seeds were lysed in Wheat Germ Lysis Buffer: 20 mM HEPES pH 7.6, 130 mM K-acetate, 5 mM Mg acetate, 0.1 mM EDTA, 10 % glycerol, 1 mM DTT, protease and phosphatase inhibitors as above. In some cases, the crude homogenate was prefiltered through Miracloth (Millipore) to remove debris, or 14,000 × g supernatant required additional clarifying centrifugation of 10 minutes at 40,000 × g.

#### Nuclear extracts

Nuclei were prepared from fresh broccoli using the CelLytic PN kit (Sigma-Aldrich). Briefly, fresh floret tips were shaved using a sharp knife into a beaker on ice and homogenized in a chilled mortar and pestle on ice in the Nuclear Isolation Buffer provided. The homogenate was further processed by brief treatment on ice with a Tissue Tearor (BioSpec Products, Inc.). Manufacturer’s instructions were followed for lysis using 1.0 % Triton X-100 and the “Semi-pure Preparation of Nuclei” protocol with 1.5 M sucrose. The resultant nuclear pellet was extracted twice with Nuclear Extraction Buffer, and the two extraction supernatants were pooled.

Coconut (*Cocos nucifera*) nuclei were purified using a composite method based on (Cutter et al., 1952; Matzke et al., 1992; Mondal et al., 1972). A young (5 month old, ~ 1.5 kg) green coconut supplied fresh from Coconut Fields Forever (Davie, FL) was opened, the liquid was drained, and the gelatinous endosperm removed by gentle scraping. Four volumes of ice-cold 4% sucrose were added to the endosperm and the mixture was homogenized 15 strokes with a loose pestle. All subsequent steps were on ice or at 4°C. The homogenate was filtered through cheesecloth to remove debris, and the nuclei were recovered from the filtrate by centrifugation at 250 × g, 10 min, 4°C. Nuclei were washed once with 15 ml Nuclear Isolation Buffer from the CelLytic PN kit (Sigma-Aldrich) and re-pelleted. After a second wash in 0.5 ml Nuclear Isolation Buffer, the recovered nuclear pellet was extracted as for broccoli nuclei using Nuclear Extraction Buffer (CelLytic PN kit, Sigma-Aldrich).

*Chlamydomonas reinhardtii* culture (2.5 liters of UTEX 90) was pelleted by centrifugation 10 minutes at 1,100 × g with no brake and washed with 900 ml 10 mM HEPES pH 7.6 at room temperature. Washed cells were collected by centrifugation 10 minutes 1,800 × g room temperature with no brake and the pellet was frozen in liquid nitrogen, thawed, refrozen and ground in a liquid nitrogen-chilled mortar and pestle. Powder was thawed and refrozen for a second grinding. After resuspension in Plant Lysis Buffer, cells still appeared largely intact so the homogenate was sonicated on ice with a probe tip 6 × 10 sec, amplitude 28%, and 6 × 10 sec amplitude 70%. This lysate was incubated 30 minutes 4°C rotating end-over-end with subsequent steps as for other green tissues.

### Fern transcriptome sequencing

As *Ceratopteris richardii* currently lacks an annotated genome, we *de novo* assembled the fern transcriptome from mRNA sequencing data collected from fronds, mature gametophytes, and spores as follows:

#### RNA sequencing

Spores were collected from adult plants and cultured on agar to gametophyte stage. Fronds were cut from healthy adult plants and flash-frozen in liquid nitrogen. Both fronds and gametophytes were ground to a fine powder prior to RNA extraction. Total RNA was extracted from spores by phenol-chloroform extraction, as previously described (Salmi and Roux, 2008). Total RNA was extracted from gametophytes, and fronds with the Spectrum Plant Total RNA Kit (Sigma-Aldrich) according to manufacturer’s instructions. Total RNA from each sample was diluted to 10-100 ng/µl and further prepared and sequenced by the Genome Sequence and Analysis Facility at UT Austin as follows: Total RNA was Poly-A-selected using the Poly(A)Purist Magnetic Kit (Life Technologies AM1919) and fragmented to ~200 bp. First strand synthesis was performed using the NEBNext First Strand Synthesis Module (NEB E7525L) with random primers and reverse transcriptase in the presence of Actinomycin D and Murine RNase Inhibitor. Second strand synthesis was performed using the NEBNExt Ultra Directional RNA Second Strand Synthesis Module (NEB E7750L) for 1 hour 16°C, and DNA was purified on AMPURE XP Beads (Beckman Coulter A63881). Double-stranded DNA was subjected to end repair and dA-tailing using the NEBNext End Repair and dA-Tailing Modules (NEB E6050L and E6053L) followed by ligation of barcode adapters (IDT) with T4Ligase (NEB). Following PCR amplification, the quality of the final library was confirmed by Bioanalyzer (Agilent). We collected 2×150 bp RNA-seq reads from fern spore, mature gametophyte, and adult frond on an Illumina HiSeq 4000 with a target of 30 million reads per tissue.

#### Fern transcriptome assembly

After removing low-quality reads (reads lacking all four nucleotides or with a no-call), we assembled transcripts with Velvet (version 1.2.06) and Oases (version 0.2.06) using each of five k-mer values (k=45, 55, 65, 75, 85). Also, we converted .fastq file to a non-redundant .fasta file and performed separate *de novo* transcriptome assembly with k=35, 45, 55, 65, and 75. Assembled transcripts were combined for each tissue, then redundant or fragmented sequences were removed based on BLASTN analysis. We determined the translational reading frame and corresponding peptide sequences from each assembled transcript based on BLASTP mapping results (after 6-frame translation *in silico*) to four plant reference proteome databases (Creinhardtii_169, Osativa_193_pep, Smoellendorffii_91_pep, TAIR10). Sequences lacking significant BLASTP scores to the reference proteomes were considered to be non-coding and omitted from the resulting fern proteome database. The resulting protein sequences derived from the three tissues were combined and a non-redundant protein sequence set computed based on clustering with UCLUST (version 4.2.66), requiring >97% amino acid identity. The supporting code is available from the NuevoTx repository (https://github.com/taejoonlab/NuevoTx). Fern transcriptome and proteome assembly statistics are summarized in **Table S4**.

### HPLC chromatography

Lysates were fractionated on a Dionex UltiMate3000 HPLC system consisting of an RS pump, Diode Array Detector, PCM-3000 pH and Conductivity Monitor, Automated Fraction Collector (Thermo Scientific, CA, USA) and a Rheodyne MXII Valve (IDEX Health & Science LLC, Rohnert Park, CA) using biocompatible PEEK tubing and either size exclusion chromatography or one of two ion exchange separations (mixed bed, or triple-phase WAXWAXCAT, see below). The sample loaded was 1-4 mg protein as measured by the BioRad Protein Assay or DC Protein Assay as appropriate to the sample buffer. Fractions were collected into 96-deep well plates. Select support ribs in the base were notched with a single-edged razor blade prior to fraction collection to accommodate subsequent use of the Life Technologies magnetic plate for bead-based mass spec preparation of samples.

#### Size exclusion

BioSep-SEC-s4000 600 × 7.8 mm ID, particle diameter 5 µm, pore diameter 500 Å (Phenomenex, Torrance, CA). Unless otherwise specified sample was 200 µl, flow rate 0.5 ml/min, with fraction collection every 45 seconds, and mobile phase 50 mM Tris-HCl pH 7.5, 50 mM NaCl. For the maize experiment the mobile phase was PBS pH 7.2 (Gibco) and for all nuclear extract experiments, Buffer S-NE (20 mM HEPES-KOH pH 7.5, 300 mM KCl, 1.5 mM MgCl_2_, and 0.2 mM EDTA). For molecular weight estimation, molecular weight standards (Sigma-Aldrich, MWGF1000, 2-5 ug each of Thyroglobulin (T9145), β-amylase (A8781), and bovine serum albumin (A8531)) were added to 1.2 mg soy extract prior to fractionation with Buffer S-NE as mobile phase.

#### Mixed bed ion exchange

Poly CATWAX A (PolyLC Mixed-Bed WAX-WCX) 200 × 4.6 mm ID, Particle diameter 5 µm, pore diameter 100 Å (PolyLC Inc., Columbia, MD). The bed contains the cation-exchange (PolyCAT A) and anion-exchange (PolyWAX LP) materials in equal amounts. A 200-250 µl sample was loaded at ≤ 40 mM NaCl, and eluted with a 1-hour salt gradient at 0.5 ml/min with collection of 0.5 ml fractions. Gradient elution was performed with Buffer A (10 mM Tris-HCl pH 7.5, 5% glycerol, 0.01% NaN_3_), and 0-70% Buffer B (1.5 M NaCl in Buffer A).

#### Triple phase ion exchange

Three columns, each 200 × 4.6 mm ID, particle diameter 5 µm, pore diameter 100 Å, were connected in series in the following order: two PolyWAX LP columns followed by a single PolyCAT A (PolyLC, Inc, Columbia, MD). Loading, buffers, and fraction collection were as for mixed bed ion exchange above with slight modifications in flow rate and elution from the methods of (Havugimana et al., 2012). The flow rate was either 0.25 ml/min with a 120 min gradient from 5-100 %B. For the separation of nuclear extracts, the gradient was modified to a 115-minute multiphasic elution from 5-100% Buffer B.

### Isoelectric focusing

Isoelectric focusing was performed using the Agilent 3100 OFFGEL Fractionator (Agilent Technologies, Santa Clara, CA) according to the manufacturer’s application note for native protein separation (Babu CV and Palaniswamy, 2014). Samples were separated into 24 fractions using the 24 cm IPG strips pH 3-10 NL (GE HealthCare, Chicago, IL). To achieve the required low salt concentration 2.5 mg wheat germ extract was diluted with water and centrifuged 10 min., 10,000 × g 4°C prior to dilution with OFFGEL Stock Solution. Broccoli nuclear extract was dialyzed in a Slide-A-Lyzer Cassette G2 with a 10,000 MWCO (Thermo Scientific, CA, USA) against 2 changes, 2 hours each, of 5.0 mM HEPES-KOH pH 7.5, 25 mM NaCl, 0.2 mM DTT followed by one change of 0.5 mM HEPES-KOH pH 7.9 overnight. A substantial precipitate formed so the solution was clarified by centrifugation 12,000 × g 10 min, 10°C, and the soluble portion further concentrated in an Amicon Ultra filter unit with a 10 kD MWCO (MilliporeSigma, Burlington, MA). BioRad Protein Assay determined that this soluble portion retained 1.5 mg (~ 35% of the predialysis amount) which was diluted with OFFGEL Stock Solution for focusing.

### Mass spectrometry sample preparation

Samples were prepared for mass spectrometry in 96-well plate format using either ultrafiltration and in-solution digestion or SP3 bead-binding of proteins and on-bead digestion ((Hughes et al., 2014) with modifications as described below). Plates were sealed with transparent film during incubation steps.

<u>Ultrafiltration</u> was performed with an AcroPrep Advance 96-filter plate,3k MWCO (Pall) using a vacuum manifold (QIAvac 96 or Multiwell, Qiagen) at −0.75 Bar. Prior to the application of the samples, preservatives were removed from the membranes by sequential filtration of 100 µl LC/MS quality water and 100µl trypsin digestion buffer (50 mM Tris-HCl, pH 8.0, 2 mM CaCl*_2_*). Samples were concentrated to 100 µl, diluted 2 fold with trypsin digestion buffer, and concentrated again to a final volume of 50-100 µl before being transferred back to a 96-deep well plate. 50 µl 2,2,2,-trifluoroethanol (TFE) was added and samples were reduced with TCEP (Bond-Breaker, Thermo,) at a final concentration of 5 mM for 30 min. 37°C. Iodoacetamide was added to 15 mM and plates were incubated in the dark 30 min. at room temperature. Alkylation was quenched by the addition of DTT to 7.5 mM. TFE was diluted to <5% by the addition of trypsin digestion buffer. 1 µg of trypsin was added to each fraction and the sealed plate was incubated 37°C overnight. Digestion was stopped by bringing samples to 0.1% formic acid, and peptides were desalted using a 5-7 µl C18 Filter Plate (Glygen Corp) with a vacuum manifold, dried, and resuspended for mass spectrometry in 3% acetonitrile, 0.1% formic acid.

<u>Bead-based</u> sample preparation was limited primarily to SEC fractionation samples due to interference of salt concentrations greater than 300 mM. Beads used were an equal mixture (vol:vol) of SpeedBead Magnetic Carboxylate Modified Particles #45152105050250 and #65152105050250 (GE Healthcare, UK). Fractions were first adjusted to 20% TFE and reduced with 5mM TCEP, 45 min, 37°C. Samples were alkylated by the addition of Iodoacetamide to 25mM final concentration, 30 minutes in the dark at room temperature, and then the reaction was quenched with 15 mM DTT. 4 µl of the mixed bead suspension (5µg/µl of each bead type) was added to each fraction. Protein binding was initiated by the addition of a premix of formic acid (to 20% of the alkylated sample) and acetonitrile (to equal 50% of final bead incubation volume. After 30 min. at room temperature with gentle rocking, the beads were pelleted by centrifugation (1000 × g, 5 min., room temperature). The deep well plate was placed on a magnetic plate (Life technologies) and all but 300 µl of the supernatant removed and discarded. After removing the sample plate from the magnet the beads were resuspended in the remaining 300 µl and transferred to a conical bottomed 450 µl 96-well plate and placed on the magnetic plate for all wash steps.

After beads were collected, the supernatant was removed and discarded and beads were washed rapidly and sequentially with 2 × 100 ml aliquots of 70% ethanol and 2 × 100 µl aliquots of acetonitrile. The plate was removed from the magnet and beads were allowed to air dry briefly before resuspension in 25 ml of 10% TFE/90% trypsin digestion buffer. 25 µl of trypsin digestion buffer containing 0.25 µg trypsin was then added to each resuspended sample for digestion overnight, 37°C. The digestion plate was placed on the magnet and the supernatant containing digested peptides was transferred to a fresh 450 µl 96-well plate and digestion stopped by bringing samples to 1% formic acid. Further peptides were recovered from the beads with two successive elution steps using 50 µl 2% DMSO and pooled with the original peptide supernatant. During the first DMSO elution, the plate was sealed and sonicated in a bath for 10 min. Peptides in the 150 µl total eluates were desalted as above.

### Chemical cross-linking

Extract (2.5 mg soy sprouts, or 2.1 mg *Chlamydomonas*) was fractionated by size exclusion as above but with PBS pH 7.2 (Gibco) as the mobile phase. DSSO (Thermo Scientific) was dissolved immediately before use in dry dimethylformamide (stored under nitrogen) at a concentration of 50 mM and then further diluted in PBS for a working stock. Immediately after fractionation working stock DSSO was added to fractions to a final concentration of 0.5 mM and samples were incubated 1 hour at room temperature. Cross-linking was quenched by the addition of 1 M Tris pH 8.0 to a concentration of 24 mM. Cross-linked fractions were immediately frozen and stored −80°C until prepared for mass spectrometry using the ultrafiltration and in-solution digestion method. Soy samples were digested as usual with trypsin, while *Chlamydomonas* samples were split after reduction/alkylation and treated as follows. One set was digested with trypsin under standard conditions while the other set was diluted 8 fold with 50 mM Tris-HCl pH 8.0, 10 mM CaCl*_2_* and digested with 0.5 µg chymotrypsin overnight 37°C. Both digests were stopped by adjusting samples to 0.1% formic acid. Peptides were desalted as above and dissolved in 20µl 5% acetonitrile, 0.1% formic acid for mass spectrometry.

### Mass spectrometry data acquisition and processing

#### Acquisition

Mass spectra were acquired using one of three Thermo mass spectrometers: Orbitrap Elite, Orbitrap Fusion, or Orbitrap Fusion Lumos. In all cases, peptides were separated using reverse phase chromatography on a Dionex Ultimate 3000 RSLCnano UHPLC system (Thermo Scientific) with a C18 trap to Acclaim C18 PepMap RSLC column (Dionex; Thermo Scientific) configuration. Peptides were eluted using a 5–40% acetonitrile gradient in 0.1% formic acid over 120 min. (for Orbitrap Elite), or a 3-45% gradient over 60 min. (for Fusion and Lumos) and directly injected into the mass spectrometer using nano-electrospray for data-dependent tandem mass spectrometry.

Data acquisition methods were as described below for each machine:

*Orbitrap Elite*: top 20 CID with full precursor ion scans (MS1) collected at 60,000 m/z resolution. Monoisotopic precursor selection and charge-state screening were enabled, with ions of charge > + 1 selected for collision-induced dissociation (CID). Up to 20 fragmentation scans (MS2) were collected per MS1. Dynamic exclusion was active with 45 s exclusion for ions selected twice within a 30 s window.

*Orbitrap Fusion*: top speed CID with full precursor ion scans (MS1) collected at 120,000 m/z resolution and a cycle time of 3 sec. Monoisotopic precursor selection and charge-state screening were enabled, with ions of charge > + 1 selected for collision-induced dissociation (CID). Dynamic exclusion was active with 60 s exclusion for ions selected once within a 60 s window. For some experiments, a similar top speed method was used with dynamic exclusion of 30 s for ions selected once within a 30 s window and high energy-induced dissociation (HCD) collision energy 31% stepped +/− 4%. All MS2 scans were centroid and done in rapid mode.

*Orbitrap Lumos*: top speed HCD with full precursor ion scans (MS1) collected at 120,000 m/z resolution. Monoisotopic precursor selection and charge-state screening were enabled using Advanced Peak Determination (APD), with ions of charge > + 1 selected for high energy-induced dissociation (HCD) with collision energy 30% stepped +/− 3%. Dynamic exclusion was active with 20 s exclusion for ions selected twice within a 20s window. All MS2 scans were centroid and done in rapid mode.

For identification of DSSO cross-linked peptides, peptides were resolved using a reverse phase nanoflow chromatography system with a 115 min 3-42% acetonitrile gradient in 0.1% formic acid. The top speed method collected full precursor ion scans (MS1) in the Orbitrap at 120,000 m/z resolution for peptides of charge 4-8 and with dynamic exclusion of 60 sec after selecting once, and a cycle time of 5 sec. CID dissociation (25% energy 10 msec) of the cross-linker was followed by MS2 scans collected in the orbitrap at 30,000 m/z resolution for charge states 2-6 using an isolation window of 1.6. Peptide pairs with a targeted mass difference of 31.9721 were selected for HCD (30% energy) and collection of rapid scan rate centroid MS3 spectra in the Ion Trap.

### Computational analyses

#### Proteomes

Reference proteomes for *Arabidopsis thaliana* (ARATH)*, Brassica oleracea* (BRAOL), *Solanum lycopersicum* (SOLLC), *Glycine max* (SOYBN), *Oryza sativa var. Japonica* (ORYSJ), *Triticum aestivum* (WHEAT), *Zea mays* (MAIZE), *Selaginella moellendorffii* (SELML), *Chlamydomonas reinhardtii* (CHLRE), were downloaded from Uniprot.org (UniProt Consortium, 2019) in August 2018. The *Chenopodium quinoa* (CHQUI) proteome was the Cquinoa_392_v1.0 assembly (Jarvis et al., 2017) downloaded from https://phytozome.jgi.doe.gov (Goodstein et al., 2012), the *Cocos nucifera* (COCNU) proteome (Armero et al., 2017) was downloaded from http://palm-comparomics.southgreen.fr, and the *Cannabis sativa* (CANSA) proteome was the canSat3 Purple Kush assembly (van Bakel et al., 2011) downloaded from http://genome.ccbr.utoronto.ca). The *Ceratopteris richardii* (CERRI) proteome was assembled *de novo* as described above.

### Initial assignment of peptide mass spectra

Peptide inference was performed with MSGF+, X!Tandem, and Comet-2013020, each run with 10ppm precursor tolerance, and allowing for fixed cysteine carbamidomethylation (+57.021464) and optional methionine oxidation (+15.9949). Peptide search results were integrated with MSBlender (Kwon et al., 2011), https://github.com/marcottelab/msblender, https://github.com/marcottelab/run_msblender). For DSSO cross-linked experiments, inter-protein cross-links were identified using the XlinkX (Klykov et al., 2018) node in ProteomeDiscover 2.2 (ThermoScientific).

### Orthogroup assignment

Each plant proteome was searched against EggNOG viridiplantae (virNOG) orthogroup HMMs using eggNOG**-**mapper v1 (Huerta-Cepas et al., 2017). Additionally, human and *Arabidopsis* proteomes were searched with the eggNOG-mapper against eukaryote (euNOG) orthogroup HMMs to allow annotation transfer of known human protein complexes. The set of proteins from all species assigned to the same orthogroup HMM were considered to belong to the same orthogroup.

### Analysis of peptide-spectral matches at the level of orthogroups

All proteomes were *in silico* trypsin digested to peptides, allowing for two missed cleavages and a missed cleavage when lysine or arginine are followed by proline. Within each proteome, only peptides from proteins within the same orthogroup were retained, and peptides matching proteins in multiple orthogroups discarded. This gave a list of orthogroup-unique peptides sometimes deriving from multiple proteins.

Experimentally-observed peptides were compared to these orthogroup unique peptides with allowance for leucine/isoleucine ambiguity to identify orthogroups present in each biochemical fraction. Peptide spectral matches (PSMS) of peptides in the same orthogroup were summed to get orthogroup PSM counts.

### Analysis of overall trends in protein recovery

We observed in at least one experiment 96.7% of the 11,339 *Viridiplantae* orthogroups that are most highly conserved *i.e.* conserved across at least half (7) of our 13 plant species. Of the 378 unobserved yet conserved orthogroups, 170 are membrane proteins suggesting their lack of detection may stem from non-ideal solubilization conditions for this class of proteins. The high coverage of conserved proteins was in marked contrast to the remaining 22,144 less conserved orthogroups, of which we only observed 58.4%. Observed proteins also tend to be longer than unobserved proteins, with a median length of 383 vs. 196 amino acids. Our observations mirror those for the extensively studied human proteome where notably 1,482/20,055 proteins have thus far evaded detection by all available proteomics technologies (Baker et al., 2017). We observed consistent tissue-specific functional trends among the observed proteins (**Figure S1**), supporting the idea that the not-yet-observed proteins, whether in plants or humans, may tend to exhibit strongly tissue or temporal specific expression or universally low abundance. Orthogroup abundances were highly reproducible across experiments (**Figure S2**).

### Assembly of features for scoring putative protein interactions

For each experiment, we assembled an elution matrix of orthogroups by fractions with PSMs normalized to 1 within each orthogroup. We additionally concatenated our training set of 32 experiments into sets by species and taxonomic group, *i.e.* all *Arabidopsis* experiments joined into one matrix and likewise for all eudicot, monocot, angiosperm, vascular plant, and green plant experiments. Maize was withheld from these concatenations as a holdout species. An elution matrix of all experiments was also made (**File S3**). Cross-linked soy and cross-linked *Chlamydomonas* experiments were not used in training. We retained only orthogroups with at least 32 total PSMs observed across the 32 combined training experiments.

We next calculated a series of all-by-all pairwise scores between orthogroup elution profiles for all 46 training matrices - Pearson’s r, Spearman’s rho, Euclidean distance, Bray-Curtis similarity, and stationary cross-correlation, all with added Poisson noise. Euclidean distance and Bray-Curtis similarity scores were inverted and normalized to a max score of 1. We calculated a hypergeometric score for the co-occurrence of proteins in fractions with repeated sampling of fractions (Drew et al., 2017). Prior to building a feature matrix of these scores for machine learning, we removed orthogroup pairs that did not correlate with at least a Pearson r > 0.3 in at least three experiments in at least two species. All scores were joined to a 3,076,998 row feature matrix of orthogroup-orthogroup similarity scores, and missing values filled with zeros.

### Construction of the gold standard protein complex training and test sets

We used known human protein complexes from the CORUM database (Giurgiu et al., 2019) as a gold standard set of positive stable protein-protein interactions. Human protein identifiers were converted to virNOG orthogroup identifiers *via* orthology to *Arabidopsis* proteins. 397 CORUM protein complexes were supplemented with 6 well-characterized plant protein complexes. Known complexes were divided into positive training and test complexes according to the scheme from (Drew et al., 2017) and complexes with over 30 members removed. We additionally supplemented 8,662 CORUM pairwise interactions with 2,562 pairs of plant proteins with direct evidence of stable protein-protein interaction, e.g. co-crystallization, co-immunoprecipitation, collected from the TAIR (Swarbreck et al., 2008) protein-protein interactions repository (ftp://ftp.arabidopsis.org/home/tair/Proteins). 20,000 negative training and test interactions were drawn from feature matrix rows, removing any interactions present in the positive gold standard set. 1240 positive training interactions and 884 positive test interactions were present in the feature matrix.

### Identification of interacting proteins by supervised machine learning

We first used the scikit-learn ExtraTreesClassifier feature selection to reduce the dimensionality of the feature matrix to the top 100 features based on declining feature importance (**Figure S3**). We used the TPOT (Olson and Moore, 2016) AutoML wrapper of scikit-learn machine learning functions for all subsequent training steps. We discovered optimal hyperparameters for an ExtraTreesClassifier with 5-fold cross-validation, with an area under the precision-recall curve of 0.64. We then trained an Extra Trees Classifier with TPOT discovered hyperparameters. This model was applied to the entire feature matrix to give a Co-Fractionation Mass Spectrometry (CF-MS) score to each pair of orthogroups, with higher scores corresponding to higher co-elution. Precision, recall, and false discovery rates were calculated from 886 positive and 20,000 negative test set interactions.

### Clustering of interacting protein pairs to define multiprotein assemblies

Interaction scores above a 10% false discovery rate threshold (CF-MS score ≥ 0.509) were input into R igraph cluster_walktrap to define coherent protein complexes. Walktrap reweighted edges between orthogroups were reformatted to a dendrogram and cut at intervals to obtain a nested hierarchy of complexes. Cuts closer to the root of the dendrogram result in larger complexes and cuts closer to the tips further define subcomplexes. A portion of these clusters are homodimers or heterodimers of closely related proteins, such as two inactive GDSL proteins, two ferredoxins, or two NTF2 proteins (**Table S5**).

### Size calibration

Based on known molecular weight size standards spiked into one soy size exclusion experiment, we fit a linear model of log10 molecular weight ~ fraction number. To transfer this calibration to other size exclusion experiments, we selected a series of internal standards complexes and proteins with known native molecular weights and consistent single peak elution patterns. A model derived from molecular weight standard spike-ins of 667, 443, 200 and 66 kDa was able to predict molecular weight values for our internal standards close to their known elution positions (**Figure S4**). We fit a new linear model for derived weights of internal standards and applied this model to all size exclusion experiments to obtain a molecular weight for each fraction.

### External datasets

For comparison purposes, known plant protein-protein interactions were downloaded from the HitPredict database (López et al., 2015) and mRNA co-expression linkages from AraNet and RiceNet (Lee et al., 2010, 2011). RNA expression Transcripts per Million (TPM) for *Chlamydomonas* and *Arabidopsis* were downloaded from the Expression Atlas (Papatheodorou et al., 2018), experiment codes E-GEOD-62671, E-GEOD-38612, E-GEOD-55866, and E-GEOD-30720. Loss-of-function annotations were assembled from (Lloyd and Meinke, 2012), Uniprot, and TAIR, and phenotype ontology obtained from the Plant PhenomeNET project (Oellrich et al., 2015). Additionally, *Arabidopsis* protein molecular weights, GO annotations, functions, unipathway, BioCyc, Reactome, BRENDA, enzyme commission, and tissue were downloaded from Uniprot as annotations to guide interpretation.

### Protein Parts Per Million (PPM) calculation

We calculated protein parts per million (PPM) following (Weiss et al., 2010), with scripts stored at github.com/marcottelab/MS_grouped_lookup/ppm_utils. Briefly, unique tryptic peptides were filtered to peptides between 7 and 40 amino acids long, and a correction factor calculated from the sum of the total length of peptides in this range per orthogroup. Observed peptide PSMs were multiplied by the peptide length, summed by orthogroup, divided by the orthogroup correction factor, multiplied by 1,000,000 and divided by the experiment total to get parts per million.

### 3D homology modeling

Precomputed 3D homology models from SWISS-MODEL (Bienert et al., 2017) were used to evaluate consistency with cross-linking data as follows: SWISS-MODEL 3D models of *Arabidopsis* CCT subunits were aligned to the known experimental *Saccharomyces cerevisiae* CCT molecular assembly (PDB: 4V94), then soy CCT subunit cross-links positioned as guided by sequence alignment to the corresponding *Arabidopsis* subunits. *Chlamydomonas* cross-links were evaluated using SWISS-MODEL 3D models of *Chlamydomonas* Photosystem II subunits that were aligned to the *Arabidopsis* Photosystem II (PDB: 5MDX). 3D models were visualized using Chimera (Pettersen et al., 2004).

### Arabidopsis phenotyping

#### Growth conditions

T-DNA mutant insertion line seeds from the SALK T-DNA insertion collection (O’Malley et al., 2015) were obtained from the Arabidopsis Biological Resource Center (https://abrc.osu.edu), and wild-type Col-0 seeds were provided by the Z.J. Chen lab. Purchased strains were SALK_056025 (*3βhsd/d2*, AT2G26260), CS832348/SAIL_726_H02 (*vdac2*, AT5G67500), CS1002787/SK15485(*domino1*, AT5G62440), CS823259/SAIL_548_H11 (*la1*, AT4G32720).

50 *Arabidopsis* seeds per genotype were sterilized for 10 minutes in 20% Clorox bleach and washed five times with 1mL diH*_2_*O. Sterilized seeds were plated on 0.5X Murashige and Skoog agar plates, sealed with parafilm, wrapped in aluminum foil and incubated at 4°C for 3-5 days, followed by incubation in a lighted growth chamber for 7 days. Plates were germinated and plants grown under an illumination cycle of 16 hours 22°C days and 8 hour 20°C nights. Seedlings were planted in a soil mix of 75% Pro-Mix Biofungicide with wetting agent/25% Profile Field and Fairway calcined clay supplemented with 1g Miracle-Gro Plant Food/gallon water, and one teaspoon Gnatrol Biological Larvicide (Valent Biosciences LLC). Bonide copper sulfate was sprayed weekly to prevent fungus. All mutant lines except CS832348 were grown an additional generation prior to phenotyping and quantitation. Days to flower were marked from the introduction of plates to light to the first visible petal, and siliques were counted at the end of flowering.

#### Genotyping

A portion of rosette leaf from each plant was flash-frozen in Eppendorf tubes in liquid nitrogen and ground to a fine powder with sterile rods. 400µl extraction buffer (200mM Tris-Cl pH 7.5, 250mM NaCl, 25mM ETDA, 0.5% SDS) was added to each tube and mixed briefly. Tubes were centrifuged at 3000 rcf for 7 minutes, then 350µl supernatant transferred to a 96 well 2mL plate prefilled with 350µl isopropanol per well, mixed by pipette, and incubated at room temperature for 10 minutes. Following centrifuging for 35 minutes at 3000 rcf, supernatant was removed by quickly inverting the plate. 150µl ethanol was added to each well, plates pulse spun, and supernatant again poured off with quick inversion of the plate. After air-drying for at least 15 minutes DNA was resuspended in 150µl H*_2_*O.

PCR reactions to confirm the genotyping were set up as two separate reactions per mutant, one for wild type with a gene specific left primer (LP) and right primer (RP), and a second with the gene specific right primer (RP) and a left border primer for the T-DNA insert (L3 or LBb1.3), according to the instructions at http://signal.salk.edu/tdnaprimers.2.html. PCR cycle conditions were: denature 98°C, anneal 55°C, extend 72°C for 35 cycles.

#### Primers

SALK T-DNA insertion line primers:

LP_SALK_056025_3BHSD GAGAGGTTCATGCTTCGACAC

RP_SALK_056025_3BHSD AGCTGCCATATGAAACACCAC

LP_SALK_056025_500maxn TGACAATAGGTGGAGTGGTCC

RP_SALK_056025_500maxn AGCTGCCATATGAAACACCAC

LBb1.3 ATTTTGCCGATTTCGGAAC

Syngenta T-DNA insertion line primers:

LP_SAIL_726_H02_VDAC CCATCAGGAGCTAGGCCTAAC

RP_SAIL_726_H02_VDAC TAAGCAGCGCACCTAAAGAAG’

LP_SAIL_548_H11 GTCTTTGCTGGTCAGGAGTTG

RP_SAIL_548_H11 CTTCTGAGATTTGTTCCAGCG

LP_SK15485 AATCCGAATACCGAATATCGG

RP_SK15485 TAAATTGGACTCCTTTGCAGC

L3 TAGCATCTGAATTTCATAACCAATCTCGATACAC

## SUPPLEMENTAL INFORMATION

Supplemental Information includes 4 figures and 5 tables.

## SUPPLEMENTAL FIGURES

**Figure S1. Related to Figure 2.**
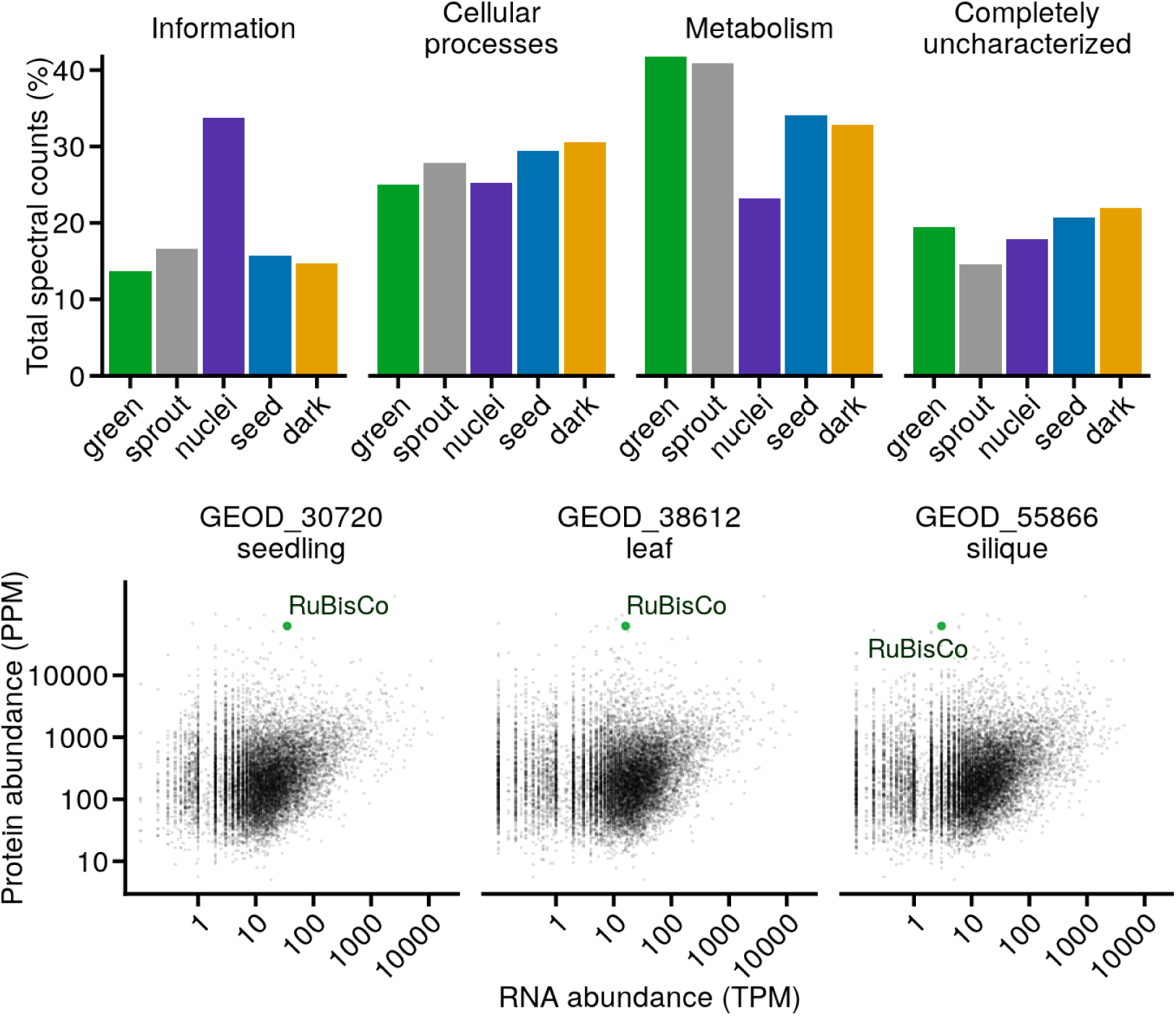
**(Top)** Tissue enrichment of COG functional categories by tissue. Proportion of total spectral counts for each tissue annotated with each high-level COG category. Nuclear samples show enrichment for information and depletion for metabolism. 15-20% of spectral counts from all tissues derive from orthogroups with no COG annotation. **(Bottom)** *Arabidopsis* mRNA vs protein abundances. RuBisCo is observed to have relatively moderate mRNA transcript levels across *Arabidopsis* tissues but is frequently the most abundant protein in our experiments. The median protein abundance of RuBisCo protein in *Arabidopsis* leaves was plotted against RNA abundance for three separate green *Arabidopsis* tissues (Gan et al., 2011; Liu et al., 2012, 2016). Titles indicate accession numbers from the ExpressionAtlas database (Petryszak et al., 2016).

**Figure S2.**
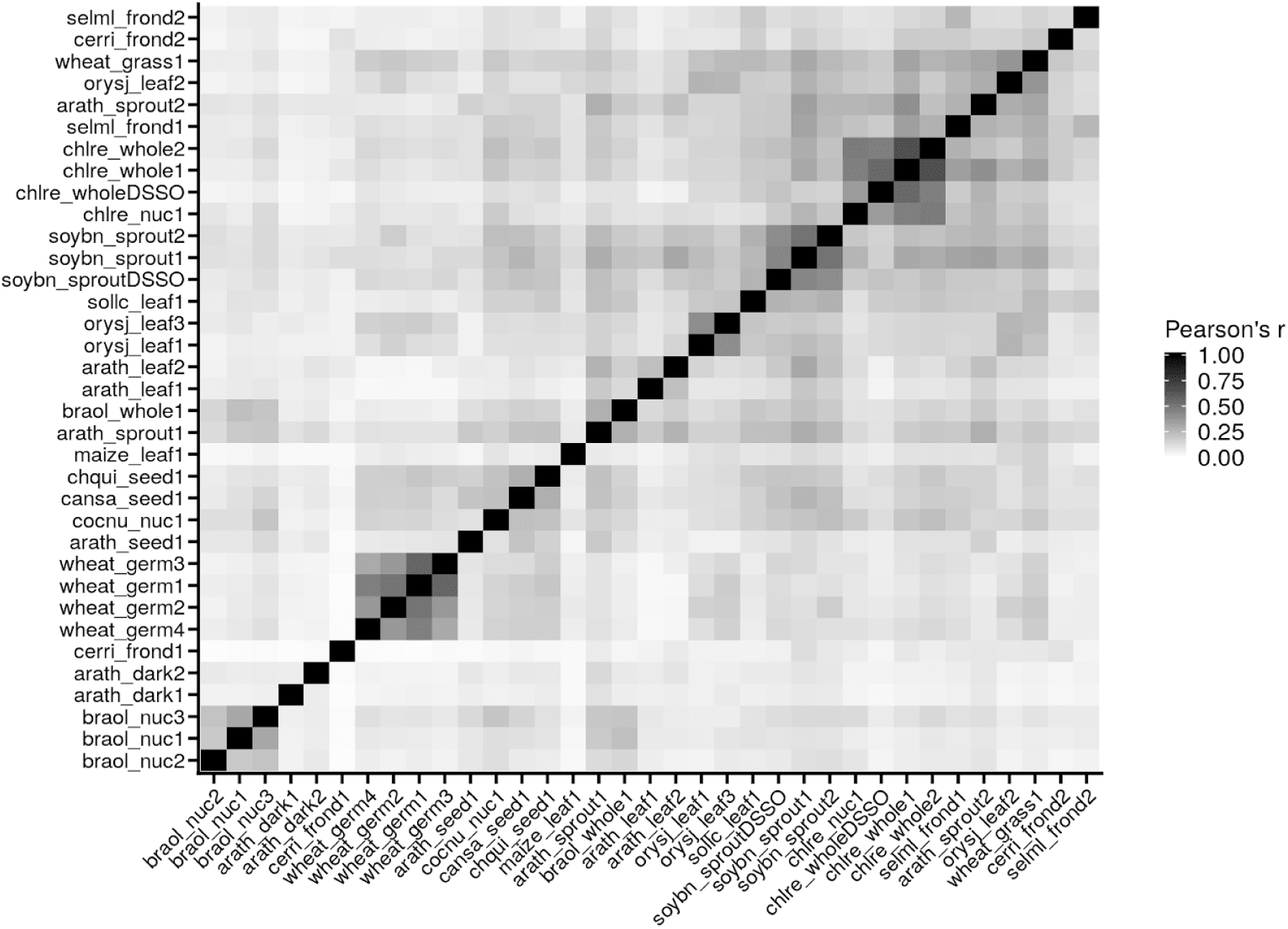
See Methods. Reproducibility of protein abundance measurements per CF-MS fractionation experiment. Hierarchically clustered all-by-all Pearson correlation of total orthogroup abundance vectors for each experiment. Experiments performed on the same tissue have highly repeatable orthogroup abundances, e.g. wheat germ and *Chlamydomonas* experiments.

**Figure S3.**
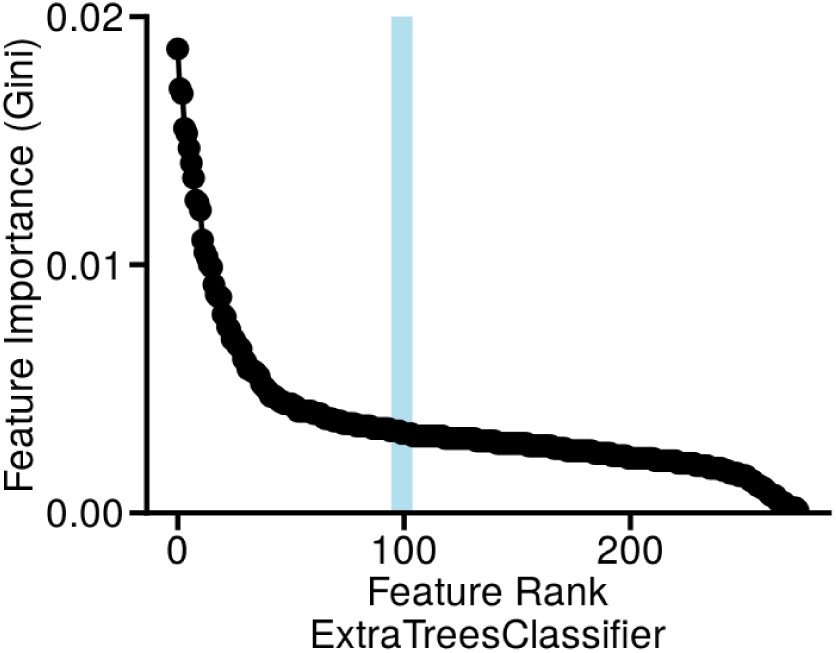
See Methods. Feature selection for protein-protein interaction scoring, as measured by scikit-learn Feature Importance. The top-ranked 100 features were selected for cross-validated model training and testing.

**Figure S4.**
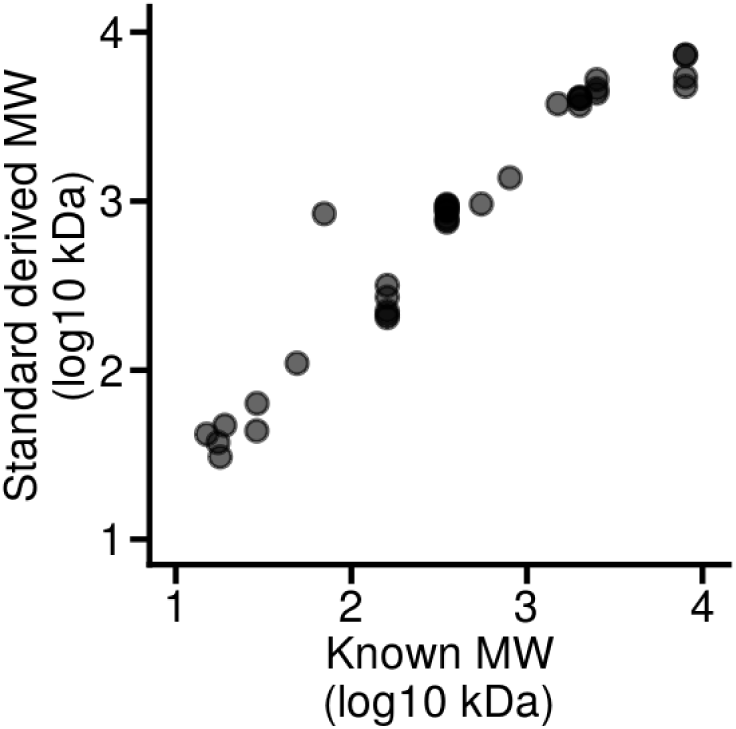
See Methods. Calibration of molecular weight estimation by size exclusion chromatography. Proteins previously known to elute at particular molecular weights in size exclusion chromatography were selected as internal reference standards, for example, members of the COP9 signalosome complex, which are known to elute at a position corresponding to approximately 350 kDa. These literature-annotated elution positions (x-axis) were compared to observed elution positions (y-axis). We observe generally excellent agreement between expected and observed molecular weights (**Spearman r = 0.97**), with one significant outlier (an Hsp70) consistently observed above 70kDa, showing that this protein participates in higher-order assemblies in plants.

## SUPPLEMENTAL TABLES

**Table S1. Related to Figure 1.**
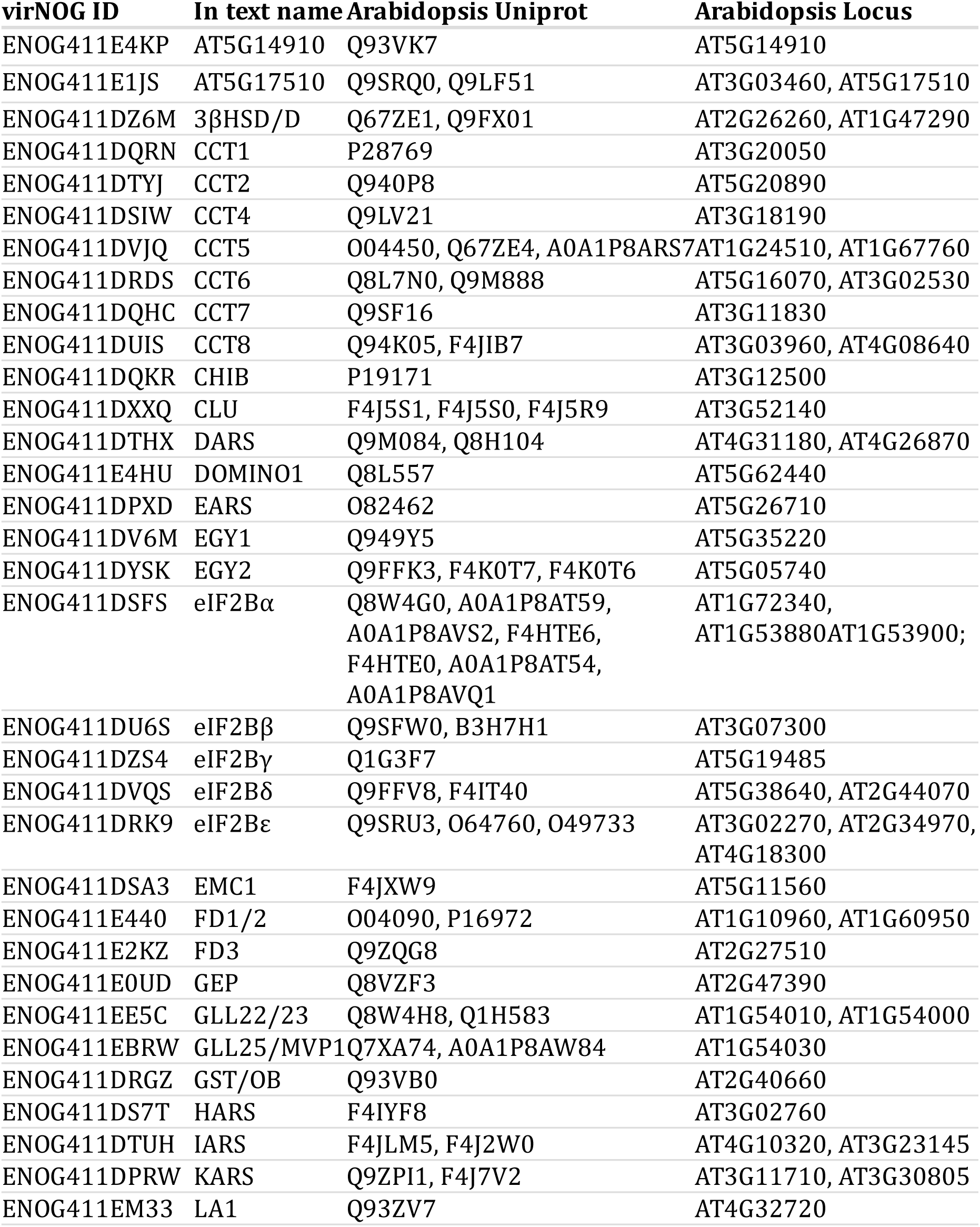
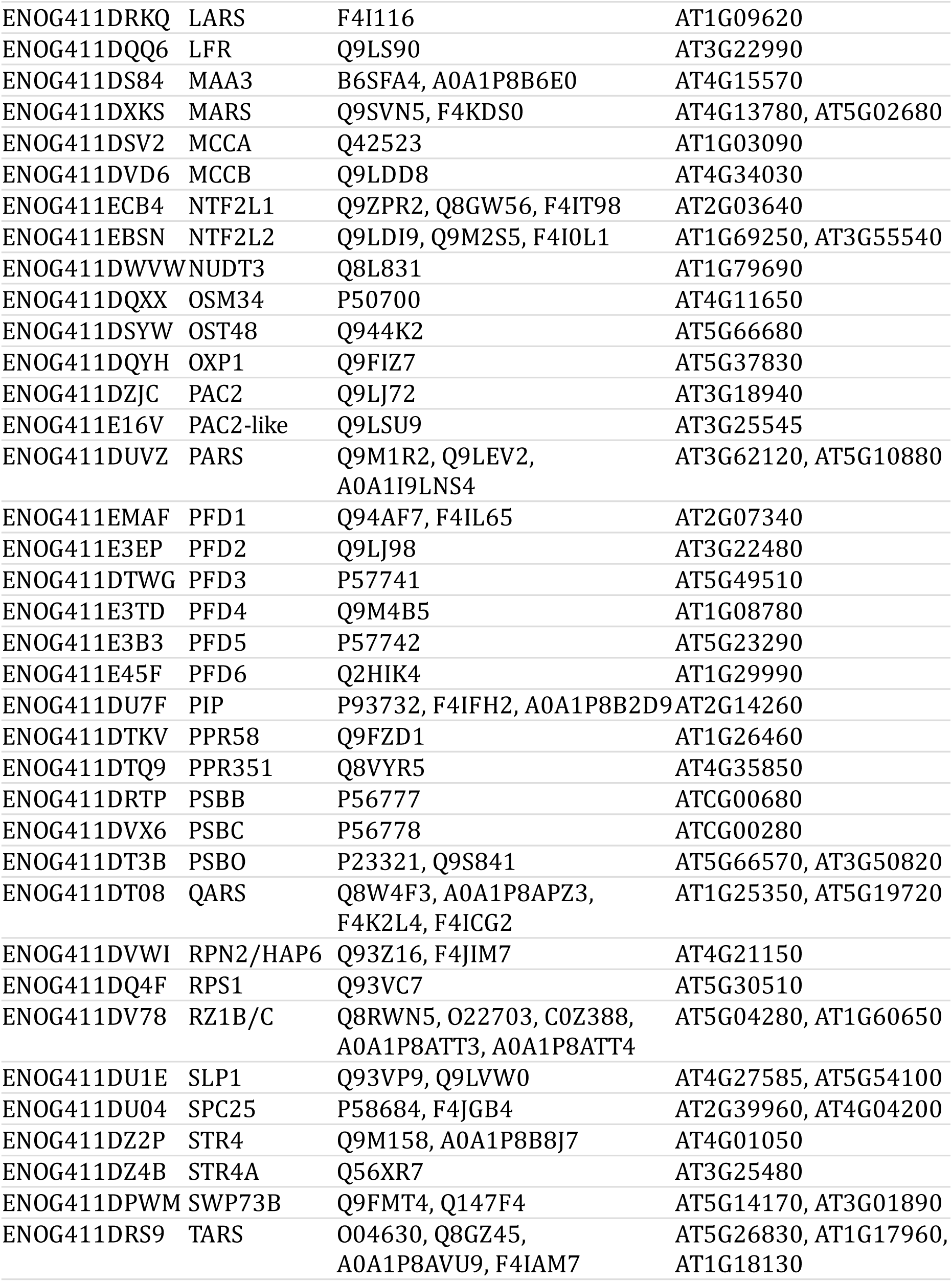
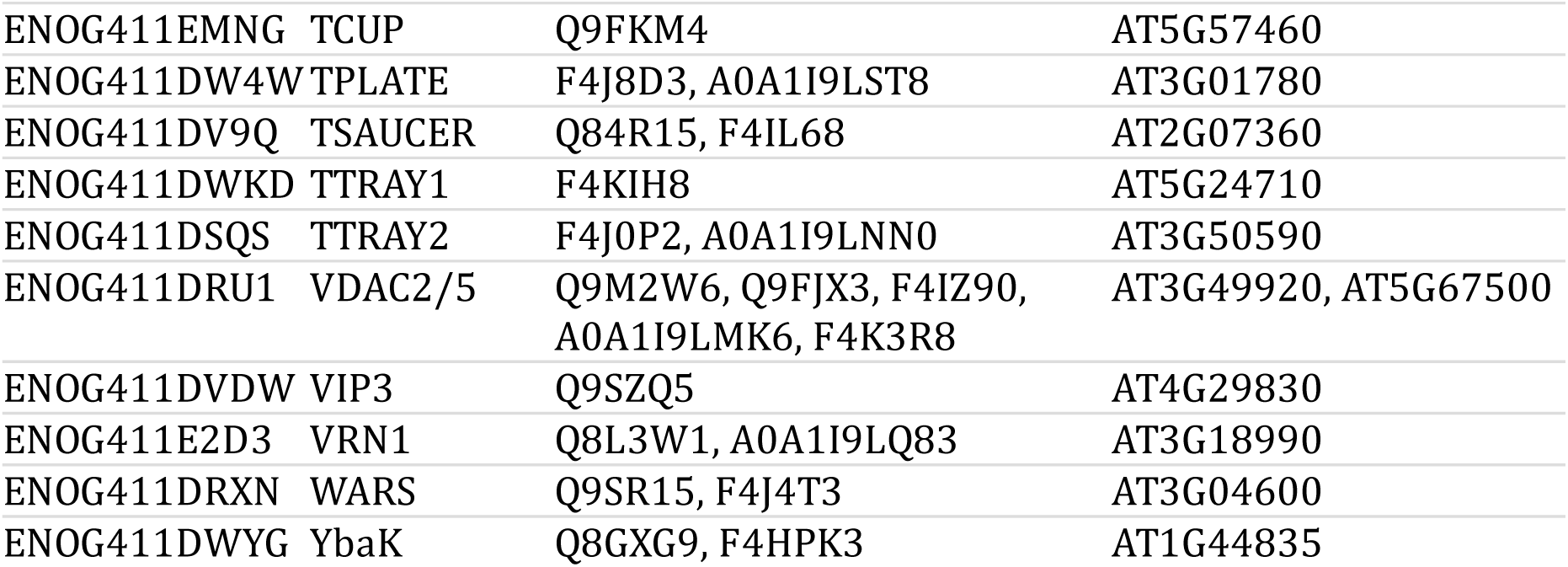
Translation of common names referenced in manuscript to OrthogroupID, *Arabidopsis* Uniprot Accession, and *Arabidopsis* locus identifiers. One orthogroup may contain multiple proteins due to duplication since the last common ancestor of plants.

**Table S2. Related to Figure 5.** The core EIF2B proteins β,δ,γ,ε are the top-scoring protein-protein interaction partners of peripheral EIF2B member EIB2Bα.

| **ID1** | **Gene1** | **ID2** | **Gene2** | **CF-MS score** |
| --- | --- | --- | --- | --- |
| ENOG411DSFS | **eIF2Bα** | ENOG411DU6S | eIF2Bβ | 0.46 |
| ENOG411DSFS | **eIF2Bα** | ENOG411DRK9 | eIF2Bε | 0.37 |
| ENOG411DSFS | **eIF2Bα** | ENOG411DZS4 | eIF2Bγ | 0.36 |
| ENOG411DSFS | **eIF2Bα** | ENOG411DVQS | eIF2Bδ | 0.35 |

**Table S3. Related to Figure 7.** Interacting partners share mutational phenotypes (here, from the plant PhenomeNET ontology (Oellrich et al., 2015)) at a significantly higher rate than random protein pairs (*p* < 10^−16^, chi-squared test).

|  | Does not share<br>phenotype | Shares<br>phenotype | Proportion |
| --- | --- | --- | --- |
| Top 10% FDR<br>interactions | 416 | 127 | 0.234 |
| Other<br>interactions | 155936 | 11326 | 0.0677 |

**Table S4.** See Methods. Assembly statistics of the *Ceratopteris richardii* non-redundant transcriptome and proteome.

|  | **Frond** | **Gametophyte** | **Spore** | **Total** |
| --- | --- | --- | --- | --- |
| # Transcripts | 369520 | 452448 | 200905 |  |
| Mean Length (bp) | 1513 | 1151 | 1267 |  |
| Median Length (bp) | 1195 | 594 | 999 |  |
| Longer than 400 bp | 265681 | 257695 | 140518 |  |
| Longer than 1 kbp | 202641 | 182536 | 100318 |  |
| Longer than 5 kbp | 8798 | 7415 | 1860 |  |
| Longer than 10 kbp | 251 | 317 | 21 |  |
| # Proteins | 168958 | 158070 | 93811 | 96677 |
| Mean Length (aa) | 394 | 388 | 385 | 263 |
| Median Length (aa) | 320 | 315 | 319 | 189 |
| Longer than 100 aa | 152195 | 141596 | 83507 | 72938 |
| Longer than 1,000 aa | 7698 | 6838 | 3748 | 1736 |
| Longer than 3,000 aa | 50 | 118 | 17 | 26 |

**Table S5.**
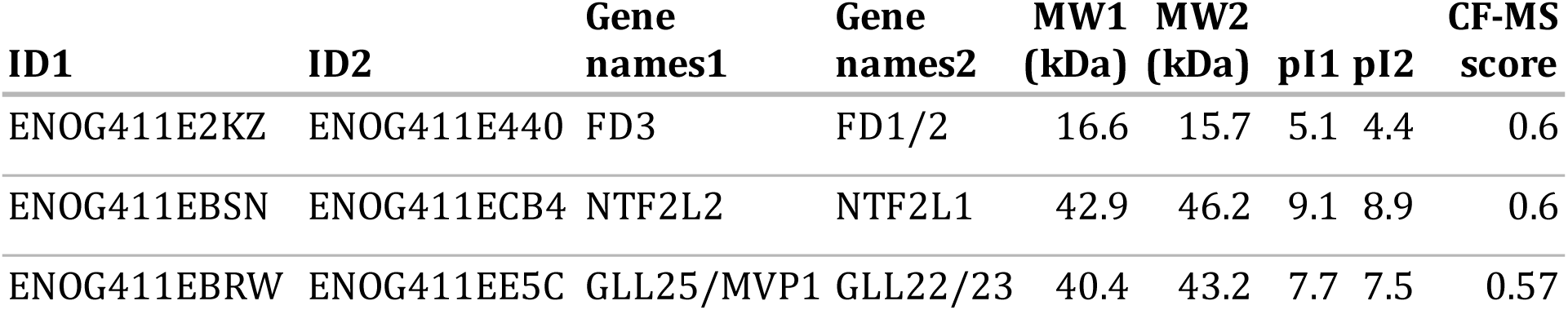
See Methods. Closely-related proteins identified as interactors. High scoring pairs of *Viridiplantae* orthogroups that belong to the same eukaryotic orthogroup. The similarity of both size and charge makes it difficult to discern whether these proteins form homomeric or heteromeric complexes.

## SUPPLEMENTARY FILES

**File S1:** Experiment attributes and observed proteins and orthogroups per fractionation *S1_experiment_numbers_plant.csv*

**File S2:** Elution matrix summarizing all fractionation data, normalized per protein per fractionation *S2_panplant_tidy_elution.csv*

**File S3:** CF-MS scores for protein-protein interactions *S3_panplant_cfms_scores_annot.txt*

**File S4:** Observed chemical cross-links in soy and *Chlamydomonas S4_crosslinks_w_score.csv*

**File S5:** Annotated walktrap clustering of interactions below FDR 10% *S5_panplant_annotated_clusters.xlsx*

## REFERENCES

Adamiec, M., Ciesielska, M., Zalaś, P., and Luciński, R. (2017). Arabidopsis thaliana intramembrane proteases. Acta Physiol. Plant. 39, 146.

Ahn, H.-K., Yoon, J.-T., Choi, I., Kim, S., Lee, H.-S., and Pai, H.-S. (2019). Functional characterization of chaperonin containing T-complex polypeptide-1 and its conserved and novel substrates in Arabidopsis. J. Exp. Bot. 70, 2741–2757.

Arabidopsis Interactome Mapping, C. (2011). Evidence for network evolution in an Arabidopsis interactome map. Science 333, 601–607.

Armero, A., Baudouin, L., Bocs, S., and This, D. (2017). Improving transcriptome de novo assembly by using a reference genome of a related species: Translational genomics from oil palm to coconut. PloS One 12, e0173300.

Aryal, U.K., McBride, Z., Chen, D., Xie, J., and Szymanski, D.B. (2017). Analysis of protein complexes in Arabidopsis leaves using size exclusion chromatography and label-free protein correlation profiling. J. Proteomics 166, 8–18.

Babu CV, S., and Palaniswamy, M.S. (2014). Agilent Application Note: Separation of Native Monoclonal Antibodies and Identification of Charge Variants.

van Bakel, H., Stout, J.M., Cote, A.G., Tallon, C.M., Sharpe, A.G., Hughes, T.R., and Page, J.E. (2011). The draft genome and transcriptome of Cannabis sativa. Genome Biol. 12, R102.

Baker, M.S., Ahn, S.B., Mohamedali, A., Islam, M.T., Cantor, D., Verhaert, P.D., Fanayan, S., Sharma, S., Nice, E.C., Connor, M., et al. (2017). Accelerating the search for the missing proteins in the human proteome. Nat. Commun. 8, 14271.

Banks, J.A., Nishiyama, T., Hasebe, M., Bowman, J.L., Gribskov, M., dePamphilis, C., Albert, V.A., Aono, N., Aoyama, T., Ambrose, B.A., et al. (2011). The Selaginella genome identifies genetic changes associated with the evolution of vascular plants. Science 332, 960–963.

Bassel, G.W., Gaudinier, A., Brady, S.M., Hennig, L., Rhee, S.Y., and De Smet, I. (2012). Systems analysis of plant functional, transcriptional, physical interaction, and metabolic networks. Plant Cell 24, 3859–3875.

Bienert, S., Waterhouse, A., de Beer, T.A.P., Tauriello, G., Studer, G., Bordoli, L., and Schwede, T. (2017). The SWISS-MODEL Repository-new features and functionality. Nucleic Acids Res. 45, D313–D319.

Browning, K.S., and Bailey-Serres, J. (2015). Mechanism of cytoplasmic mRNA translation. Arab. Book 13, e0176.

Cestari, I., Kalidas, S., Monnerat, S., Anupama, A., Phillips, M.A., and Stuart, K. (2013). A multiple aminoacyl-tRNA synthetase complex that enhances tRNA-aminoacylation in African trypanosomes. Mol. Cell. Biol. 33, 4872–4888.

Chang, R.L., Ghamsari, L., Manichaikul, A., Hom, E.F.Y., Balaji, S., Fu, W., Shen, Y., Hao, T., Palsson, B.Ø., Salehi-Ashtiani, K., et al. (2011). Metabolic network reconstruction of Chlamydomonas offers insight into light-driven algal metabolism. Mol. Syst. Biol. 7, 518.

Cutter, V.M., Wilson, K.S., and Dube, G.R. (1952). The isolation of living nuclei from the endosperm of Cocos nucifera. Science 115, 58–59.

Dhawan, R., Luo, H., Foerster, A.M., Abuqamar, S., Du, H.-N., Briggs, S.D., Mittelsten Scheid, O., and Mengiste, T. (2009). HISTONE MONOUBIQUITINATION1 interacts with a subunit of the mediator complex and regulates defense against necrotrophic fungal pathogens in Arabidopsis. Plant Cell 21, 1000–1019.

Dong, S., and Wang, Y. (2016). Nudix Effectors: A Common Weapon in the Arsenal of Plant Pathogens. PLoS Pathog. 12.

Drew, K., Lee, C., Huizar, R.L., Tu, F., Borgeson, B., McWhite, C.D., Ma, Y., Wallingford, J.B., and Marcotte, E.M. (2017). Integration of over 9,000 mass spectrometry experiments builds a global map of human protein complexes. Mol. Syst. Biol. 13, 932.

Eisenberg, D., Marcotte, E.M., Xenarios, I., and Yeates, T.O. (2000). Protein function in the post-genomic era. Nature 405, 823–826.

Feller, U., Anders, I., and Mae, T. (2007). Rubiscolytics: fate of Rubisco after its enzymatic function in a cell is terminated. J. Exp. Bot. 59, 1615–1624.

Fleurdépine, S., Deragon, J.-M., Devic, M., Guilleminot, J., and Bousquet-Antonelli, C. (2007). A bona fide La protein is required for embryogenesis in Arabidopsis thaliana. Nucleic Acids Res. 35, 3306–3321.

Fraser, H.B., and Plotkin, J.B. (2007). Using protein complexes to predict phenotypic effects of gene mutation. Genome Biol 8, R252.

Gadeyne, A., Sánchez-Rodrıguez, C., Vanneste, S., Di Rubbo, S., Zauber, H., Vanneste, K., Van Leene, J., De Winne, N., Eeckhout, D., Persiau, G., et al. (2014). The TPLATE adaptor complex drives clathrin-mediated endocytosis in plants. Cell 156, 691–704.

Gan, X., Stegle, O., Behr, J., Steffen, J.G., Drewe, P., Hildebrand, K.L., Lyngsoe, R., Schultheiss, S.J., Osborne, E.J., Sreedharan, V.T., et al. (2011). Multiple reference genomes and transcriptomes for Arabidopsis thaliana. Nature 477, 419–423.

Gilbert, M., and Schulze, W.X. (2019). Global Identification of Protein Complexes within the Membrane Proteome of Arabidopsis Roots Using a SEC-MS Approach. J. Proteome Res. 18, 107–119.

Giurgiu, M., Reinhard, J., Brauner, B., Dunger-Kaltenbach, I., Fobo, G., Frishman, G., Montrone, C., and Ruepp, A. (2019). CORUM: the comprehensive resource of mammalian protein complexes-2019. Nucleic Acids Res. 47, D559–D563.

Goodstein, D.M., Shu, S., Howson, R., Neupane, R., Hayes, R.D., Fazo, J., Mitros, T., Dirks, W., Hellsten, U., Putnam, N., et al. (2012). Phytozome: a comparative platform for green plant genomics. Nucleic Acids Res. 40, D1178–D1186.

Hartwell, L.H., Hopfield, J.J., Leibler, S., and Murray, A.W. (1999). From molecular to modular cell biology. Nature 402, C47–52.

Havugimana, P.C., Hart, G.T., Nepusz, T., Yang, H., Turinsky, A.L., Li, Z., Wang, P.I., Boutz, D.R., Fong, V., Phanse, S., et al. (2012). A census of human soluble protein complexes. Cell 150, 1068–1081.

Hein, M.Y., Hubner, N.C., Poser, I., Cox, J., Nagaraj, N., Toyoda, Y., Gak, I.A., Weisswange, I., Mansfeld, J., Buchholz, F., et al. (2015). A Human Interactome in Three Quantitative Dimensions Organized by Stoichiometries and Abundances. Cell 163, 712–723.

Hirano, Y., Hendil, K.B., Yashiroda, H., Iemura, S., Nagane, R., Hioki, Y., Natsume, T., Tanaka, K., and Murata, S. (2005). A heterodimeric complex that promotes the assembly of mammalian 20S proteasomes. Nature 437, 1381–1385.

Hu, P., Janga, S.C., Babu, M., Díaz-Mejía, J.J., Butland, G., Yang, W., Pogoutse, O., Guo, X., Phanse, S., Wong, P., et al. (2009). Global functional atlas of Escherichia coli encompassing previously uncharacterized proteins. PLoS Biol. 7, e96.

Huerta-Cepas, J., Forslund, K., Coelho, L.P., Szklarczyk, D., Jensen, L.J., von Mering, C., and Bork, P. (2017). Fast Genome-Wide Functional Annotation through Orthology Assignment by eggNOG-Mapper. Mol. Biol. Evol. 34, 2115–2122.

Hughes, C.S., Foehr, S., Garfield, D.A., Furlong, E.E., Steinmetz, L.M., and Krijgsveld, J. (2014). Ultrasensitive proteome analysis using paramagnetic bead technology. Mol. Syst. Biol. 10, 757.

Huttlin, E.L., Ting, L., Bruckner, R.J., Gebreab, F., Gygi, M.P., Szpyt, J., Tam, S., Zarraga, G., Colby, G., Baltier, K., et al. (2015). The BioPlex Network: A Systematic Exploration of the Human Interactome. Cell 162, 425–440.

Huttlin, E.L., Bruckner, R.J., Paulo, J.A., Cannon, J.R., Ting, L., Baltier, K., Colby, G., Gebreab, F., Gygi, M.P., Parzen, H., et al. (2017). Architecture of the human interactome defines protein communities and disease networks. Nature 545, 505–509.

Jarvis, D.E., Ho, Y.S., Lightfoot, D.J., Schmöckel, S.M., Li, B., Borm, T.J.A., Ohyanagi, H., Mineta, K., Michell, C.T., Saber, N., et al. (2017). The genome of Chenopodium quinoa. Nature 542, 307–312.

Jiao, Y., Wickett, N.J., Ayyampalayam, S., Chanderbali, A.S., Landherr, L., Ralph, P.E., Tomsho, L.P., Hu, Y., Liang, H., Soltis, P.S., et al. (2011). Ancestral polyploidy in seed plants and angiosperms. Nature 473, 97–100.

Kan, J., An, L., Wu, Y., Long, J., Song, L., Fang, R., and Jia, Y. (2018). A dual role for proline iminopeptidase in the regulation of bacterial motility and host immunity. Mol. Plant Pathol.

Kawahara, Y., de la Bastide, M., Hamilton, J.P., Kanamori, H., McCombie, W.R., Ouyang, S., Schwartz, D.C., Tanaka, T., Wu, J., Zhou, S., et al. (2013). Improvement of the Oryza sativa Nipponbare reference genome using next generation sequence and optical map data. Rice 6, 4.

Kirkwood, K.J., Ahmad, Y., Larance, M., and Lamond, A.I. (2013). Characterization of native protein complexes and protein isoform variation using size-fractionation-based quantitative proteomics. Mol Cell Proteomics 12, 3851–3873.

Klykov, O., Steigenberger, B., Pektas, S., Fasci, D., Heck, A.J.R., and Scheltema, R.A. (2018). Efficient and robust proteome-wide approaches for cross-linking mass spectrometry. Nat Protoc 13, 2964–2990.

Koornneef, M. (1981). The complex syndrome of ttg mutants. Arab. Inf. Serv. 18, 45–51.

Kriechbaumer, V., Maneta-Peyret, L., Fouillen, L., Botchway, S.W., Upson, J., Hughes, L., Richardson, J., Kittelmann, M., Moreau, P., and Hawes, C. (2018). The odd one out: Arabidopsis reticulon 20 does not bend ER membranes but has a role in lipid regulation. Sci. Rep. 8, 2310.

Kristensen, A.R., Gsponer, J., and Foster, L.J. (2012). A high-throughput approach for measuring temporal changes in the interactome. Nat Methods 9, 907–909.

Kumar, S., Stecher, G., Suleski, M., and Hedges, S.B. (2017). TimeTree: A Resource for Timelines, Timetrees, and Divergence Times. Mol. Biol. Evol. 34, 1812–1819.

Kwon, T., Choi, H., Vogel, C., Nesvizhskii, A.I., and Marcotte, E.M. (2011). MSblender: A probabilistic approach for integrating peptide identifications from multiple database search engines. J. Proteome Res. 10, 2949–2958.

Lage, K., Karlberg, E.O., Storling, Z.M., Olason, P.I., Pedersen, A.G., Rigina, O., Hinsby, A.M., Tumer, Z., Pociot, F., Tommerup, N., et al. (2007). A human phenome-interactome network of protein complexes implicated in genetic disorders. Nat Biotechnol 25, 309–316.

Lahmy, S., Guilleminot, J., Cheng, C.-M., Bechtold, N., Albert, S., Pelletier, G., Delseny, M., and Devic, M. (2004). DOMINO1, a member of a small plant-specific gene family, encodes a protein essential for nuclear and nucleolar functions. Plant J. Cell Mol. Biol. 39, 809–820.

Laporte, D., Huot, J.L., Bader, G., Enkler, L., Senger, B., and Becker, H.D. (2014). Exploring the evolutionary diversity and assembly modes of multi-aminoacyl-tRNA synthetase complexes: Lessons from unicellular organisms. FEBS Lett. 588, 4268–4278.

Lee, I., Ambaru, B., Thakkar, P., Marcotte, E.M., and Rhee, S.Y. (2010). Rational association of genes with traits using a genome-scale gene network for Arabidopsis thaliana. Nat Biotechnol 28, 149–156.

Lee, I., Seo, Y.S., Coltrane, D., Hwang, S., Oh, T., Marcotte, E.M., and Ronald, P.C. (2011). Genetic dissection of the biotic stress response using a genome-scale gene network for rice. Proc Natl Acad Sci U A 108, 18548–18553.

Levy, Y.Y., Mesnage, S., Mylne, J.S., Gendall, A.R., and Dean, C. (2002). Multiple roles of Arabidopsis VRN1 in vernalization and flowering time control. Science 297, 243–246.

Lintala, M., Schuck, N., Thormählen, I., Jungfer, A., Weber, K.L., Weber, A.P.M., Geigenberger, P., Soll, J., Bölter, B., and Mulo, P. (2014). Arabidopsis tic62 trol mutant lacking thylakoid-bound ferredoxin-NADP+ oxidoreductase shows distinct metabolic phenotype. Mol. Plant 7, 45–57.

Liu, J., Jung, C., Xu, J., Wang, H., Deng, S., Bernad, L., Arenas-Huertero, C., and Chua, N.-H. (2012). Genome-wide analysis uncovers regulation of long intergenic noncoding RNAs in Arabidopsis. Plant Cell 24, 4333–4345.

Liu, J., Deng, S., Wang, H., Ye, J., Wu, H.-W., Sun, H.-X., and Chua, N.-H. (2016). CURLY LEAF Regulates Gene Sets Coordinating Seed Size and Lipid Biosynthesis. Plant Physiol. 171, 424–436.

Lloyd, J., and Meinke, D. (2012). A Comprehensive Dataset of Genes with a Loss-of-Function Mutant Phenotype in Arabidopsis. Plant Physiol. 158, 1115–1129.

López, Y., Nakai, K., and Patil, A. (2015). HitPredict version 4: comprehensive reliability scoring of physical protein–protein interactions from more than 100 species. Database J. Biol. Databases Curation 2015.

Majeran, W., Zybailov, B., Ytterberg, A.J., Dunsmore, J., Sun, Q., and van Wijk, K.J. (2008). Consequences of C4 Differentiation for Chloroplast Membrane Proteomes in Maize Mesophyll and Bundle Sheath Cells. Mol. Cell. Proteomics MCP 7, 1609–1638.

Malovannaya, A., Lanz, R.B., Jung, S.Y., Bulynko, Y., Le, N.T., Chan, D.W., Ding, C., Shi, Y., Yucer, N., Krenciute, G., et al. (2011). Analysis of the human endogenous coregulator complexome. Cell 145, 787–799.

Marcotte, E.M., Pellegrini, M., Thompson, M.J., Yeates, T.O., and Eisenberg, D. (1999). A combined algorithm for genome-wide prediction of protein function. Nature 402, 83–86.

Marsh, J.A., and Teichmann, S.A. (2015). Structure, Dynamics, Assembly, and Evolution of Protein Complexes. Annu. Rev. Biochem. 84, 551–575.

Matzke, A.J., Behensky, C., Weiger, T., and Matzke, M.A. (1992). A large conductance ion channel in the nuclear envelope of a higher plant cell. FEBS Lett. 302, 81–85.

McBride, Z., Chen, D., Lee, Y., Aryal, U.K., Xie, J., and Szymanski, D.B. (2019). A Label-Free Mass Spectrometry Method to Predict Endogenous Protein Complex Composition. Mol. Cell. Proteomics mcp.RA 119.001400.

McGary, K.L., Lee, I., and Marcotte, E.M. (2007). Broad network-based predictability of Saccharomyces cerevisiae gene loss-of-function phenotypes. Genome Biol 8, R258.

Mondal, H., Mandal, R.K., and Biswas, B.B. (1972). RNA polymerase from eukaryotic cells. Isolation and purification of enzymes and factors from chromatin of coconut nuclei. Eur. J. Biochem. 25, 463–470.

Mukaihara, T., Tamura, N., Murata, Y., and Iwabuchi, M. (2004). Genetic screening of Hrp type III-related pathogenicity genes controlled by the HrpB transcriptional activator in Ralstonia solanacearum. Mol. Microbiol. 54, 863–875.

Niehaus, T.D., Thamm, A.M., de Crecy-Lagard, V., and Hanson, A.D. (2015). Proteins of Unknown Biochemical Function: A Persistent Problem and a Roadmap to Help Overcome It. Plant Physiol 169, 1436–1442.

Oellrich, A., Walls, R.L., Cannon, E.K., Cannon, S.B., Cooper, L., Gardiner, J., Gkoutos, G.V., Harper, L., He, M., Hoehndorf, R., et al. (2015). An ontology approach to comparative phenomics in plants. Plant Methods 11, 10.

Oliver, S. (2000). Guilt-by-association goes global. Nature 403, 601–603.

Olson, R.S., and Moore, J.H. (2016). TPOT: A Tree-based Pipeline Optimization Tool for Automating Machine Learning. In JMLR: Workshop and Conference Proceedings, pp. 66–74.

O’Malley, R.C., Barragan, C.C., and Ecker, J.R. (2015). A user’s guide to the Arabidopsis T-DNA insertion mutant collections. Methods Mol. Biol. Clifton NJ 1284, 323–342.

Omura, Y., Nishio, Y., Takemoto, T., Ikeuchi, C., Sekine, O., Morino, K., Maeno, Y., Obata, T., Ugi, S., Maegawa, H., et al. (2009). SAFB1, an RBMX-binding protein, is a newly identified regulator of hepatic SREBP-1c gene. BMB Rep. 42, 232–237.

Panchy, N., Wu, G., Newton, L., Tsai, C.-H., Chen, J., Benning, C., Farré, E.M., and Shiu, S.-H. (2014). Prevalence, evolution, and cis-regulation of diel transcription in Chlamydomonas reinhardtii. G3 Bethesda Md 4, 2461–2471.

Papatheodorou, I., Fonseca, N.A., Keays, M., Tang, Y.A., Barrera, E., Bazant, W., Burke, M., Füllgrabe, A., Fuentes, A.M.-P., George, N., et al. (2018). Expression Atlas: gene and protein expression across multiple studies and organisms. Nucleic Acids Res. 46, D246–D251.

Perez-Riverol, Y., Csordas, A., Bai, J., Bernal-Llinares, M., Hewapathirana, S., Kundu, D.J., Inuganti, A., Griss, J., Mayer, G., Eisenacher, M., et al. (2019). The PRIDE database and related tools and resources in 2019: improving support for quantification data. Nucleic Acids Res. 47, D442–D450.

Petryszak, R., Keays, M., Tang, Y.A., Fonseca, N.A., Barrera, E., Burdett, T., Füllgrabe., Fuentes, A.M.-P., Jupp, S., Koskinen, S., et al. (2016). Expression Atlas update—an integrated database of gene and protein expression in humans, animals and plants. Nucleic Acids Res. 44, D746–D752.

Pettersen, E.F., Goddard, T.D., Huang, C.C., Couch, G.S., Greenblatt, D.M., Meng, E.C., and Ferrin, T.E. (2004). UCSF Chimera--a visualization system for exploratory research and analysis. J. Comput. Chem. 25, 1605–1612.

Prasad, M., Kaur, J., Pawlak, K.J., Bose, M., Whittal, R.M., and Bose, H.S. (2015). Mitochondria-associated endoplasmic reticulum membrane (MAM) regulates steroidogenic activity via steroidogenic acute regulatory protein (StAR)-voltage-dependent anion channel 2 (VDAC2) interaction. J. Biol. Chem. 290, 2604–2616.

Pulido, P., Zagari, N., Manavski, N., Gawronski, P., Matthes, A., Scharff, L.B., Meurer, J., and Leister, D. (2018). CHLOROPLAST RIBOSOME ASSOCIATED Supports Translation under Stress and Interacts with the Ribosomal 30S Subunit. Plant Physiol. 177, 1539–1554.

Rhee, S.Y., and Mutwil, M. (2014). Towards revealing the functions of all genes in plants. Trends Plant Sci 19, 212–221.

Rivosecchi, J., Larochelle, M., Teste, C., Grenier, F., Malapert, A., Ricci, E.P., Bernard, P., Bachand, F., and Vanoosthuyse, V. (2019). Senataxin homologue Sen1 is required for efficient termination of RNA polymerase III transcription. EMBO J. 38, e101955.

Rohila, J.S., Chen, M., Chen, S., Chen, J., Cerny, R.L., Dardick, C., Canlas, P., Fujii, H., Gribskov, M., Kanrar, S., et al. (2009). Protein-protein interactions of tandem affinity purified protein kinases from rice. PLoS One 4, e6685.

Salmi, M.L., and Roux, S.J. (2008). Gene expression changes induced by space flight in single-cells of the fern Ceratopteris richardii. Planta 229, 151–159.

Savary, S., Willocquet, L., Pethybridge, S.J., Esker, P., McRoberts, N., and Nelson, A. (2019). The global burden of pathogens and pests on major food crops. Nat. Ecol. Evol. 3, 430–439.

Schwikowski, B., Uetz, P., and Fields, S. (2000). A network of protein-protein interactions in yeast. Nat Biotechnol 18, 1257–1261.

Shikanai, T. (2016). Chloroplast NDH: A different enzyme with a structure similar to that of respiratory NADH dehydrogenase. Biochim. Biophys. Acta 1857, 1015–1022.

Strasser, R. (2016). Plant protein glycosylation. Glycobiology 26, 926–939.

Swarbreck, D., Wilks, C., Lamesch, P., Berardini, T.Z., Garcia-Hernandez, M., Foerster, H., Li, D., Meyer, T., Muller, R., Ploetz, L., et al. (2008). The Arabidopsis Information Resource (TAIR): gene structure and function annotation. Nucleic Acids Res 36, D1009–14.

Takabayashi, A., Takabayashi, S., Takahashi, K., Watanabe, M., Uchida, H., Murakami, A., Fujita, T., Ikeuchi, M., and Tanaka, A. (2017). PCoM-DB Update: A Protein Co-Migration Database for Photosynthetic Organisms. Plant Cell Physiol. 58, e10.

Tateda, C., Watanabe, K., Kusano, T., and Takahashi, Y. (2011). Molecular and genetic characterization of the gene family encoding the voltage-dependent anion channel in Arabidopsis. J. Exp. Bot. 62, 4773–4785.

UniProt Consortium (2019). UniProt: a worldwide hub of protein knowledge. Nucleic Acids Res. 47, D506–D515.

Van Leene, J., Witters, E., Inze, D., and De Jaeger, G. (2008). Boosting tandem affinity purification of plant protein complexes. Trends Plant Sci 13, 517–520.

Van Leene, J., Hollunder, J., Eeckhout, D., Persiau, G., Van De Slijke, E., Stals, H., Van Isterdael, G., Verkest, A., Neirynck, S., Buffel, Y., et al. (2010). Targeted interactomics reveals a complex core cell cycle machinery in Arabidopsis thaliana. Mol Syst Biol 6, 397.

Van Leene, J., Boruc, J., De Jaeger, G., Russinova, E., and De Veylder, L. (2011). A kaleidoscopic view of the Arabidopsis core cell cycle interactome. Trends Plant Sci 16, 141–150.

Vercruyssen, L., Verkest, A., Gonzalez, N., Heyndrickx, K.S., Eeckhout, D., Han, S.-K., Jégu, T., Archacki, R., Van Leene, J., Andriankaja, M., et al. (2014). ANGUSTIFOLIA3 binds to SWI/SNF chromatin remodeling complexes to regulate transcription during Arabidopsis leaf development. Plant Cell 26, 210–229.

Vidal, M., Cusick, M.E., and Barabasi, A.L. (2011). Interactome networks and human disease. Cell 144, 986–998.

Vogel, C., and Marcotte, E.M. (2012). Insights into the regulation of protein abundance from proteomic and transcriptomic analyses. Nat. Rev. Genet. 13, 227–232.

Walhout, A.J., and Vidal, M. (2001). Protein interaction maps for model organisms. Nat. Rev. Mol. Cell Biol. 2, 55–62.

Wan, C., Borgeson, B., Phanse, S., Tu, F., Drew, K., Clark, G., Xiong, X., Kagan, O., Kwan, J., Bezginov, A., et al. (2015). Panorama of ancient metazoan macromolecular complexes. Nature 525, 339–344.

Weiss, M., Schrimpf, S., Hengartner, M.O., Lercher, M.J., and von Mering, C. (2010). Shotgun proteomics data from multiple organisms reveals remarkable quantitative conservation of the eukaryotic core proteome. Proteomics 10, 1297–1306.

Wu, Z., Zhu, D., Lin, X., Miao, J., Gu, L., Deng, X., Yang, Q., Sun, K., Zhu, D., Cao, X., et al. (2016). RNA Binding Proteins RZ-1B and RZ-1C Play Critical Roles in Regulating Pre-mRNA Splicing and Gene Expression during Development in Arabidopsis. Plant Cell 28, 55–73.

Zhang, Y., Gao, P., and Yuan, J.S. (2010). Plant protein-protein interaction network and interactome. Curr Genomics 11, 40–46.

